# A curated dataset of great ape genome diversity

**DOI:** 10.1101/2025.02.18.638799

**Authors:** Sojung Han, Sepand Riyahi, Xin Huang, Martin Kuhlwilm

## Abstract

Studying the genetic diversity of non-human great apes is important for research questions in evolution as well as human diversity and disease. Genomic data of the three great ape clades (*Pan*, *Gorilla*, *Pongo*) has been published across multiple studies over more than one decade. However, unlike in humans, no comprehensive dataset on great ape diversity is available, due to different scopes of the original studies. Here, we present a curated dataset of 332 high coverage (≥12-fold) whole genomes, including 198 chimpanzee, 16 bonobo, 77 gorilla and 41 orangutan individuals sequenced on the Illumina platform. By integrating data from captive individuals, we contextualize them with data from wild individuals. We discuss issues with previously published data leading to removal of individuals due to low sequencing depth, missing data, or occurrence of duplicate individuals. This resource of files in CRAM and gVCF format, as well as segregating sites per clade, will allow researchers to address questions related to human and great ape evolution and diversity in a comparative manner.

## Background & Summary

Great apes have been of long-standing interest as the closest living relatives of humans. Studying their phylogenetic relationships to humans and amongst each other has been an important field of genetic and genomic research over the past few decades^1–4^, fostering investigations into human uniqueness and understanding evolution and disease^5–7^. An assembly of the chimpanzee reference genome was published soon after the human reference genome^8^, and by now high-quality reference genomes of all great ape species are available^9–11^. However, beyond the insights from single genome assemblies, the diversity within a clade is an important aspect to understand their evolution and their species-specific traits^12,13^.

While the advent of high-throughput sequencing technologies allowed characterizing the genetic makeup of many individuals, in humans on a large scale and with high quality^14^, this is not possible in great apes. Great apes are all endangered or critically endangered^15^, with rapidly shrinking habitats and a high risk of extinction in the wild in the near future, and small captive populations from a limited pool of founder individuals. Hence, a limited number of individuals is available for genomic studies, and often, access to genetic diversity is only possible through non-invasive sampling^16^, or from other degraded sources such as historical collections^17^. However, several key publications generated diversity data from all present-day great ape species (Fig. 1): chimpanzees (*Pan troglodytes*)^18,19^, bonobos (*Pan paniscus*)^18^, western gorillas (*Gorilla gorilla*)^18,20^ and eastern gorillas (*Gorilla beringei*)^18,21,22^, as well as Bornean, Sumatran and Tapanuli orangutans (*Pongo pygmaeus*, *Pongo abelii*, *Pongo tapanuliensis*)^18,23^. This is complemented by a number of genomes from mostly captive individuals across these clades, published in the context of genome assemblies^9–11,24^, trio sequencing for the estimation of mutation rates^25–27^, functional studies^28^, or large-scale studies of primate diversity^29^.

**Figure 1.**
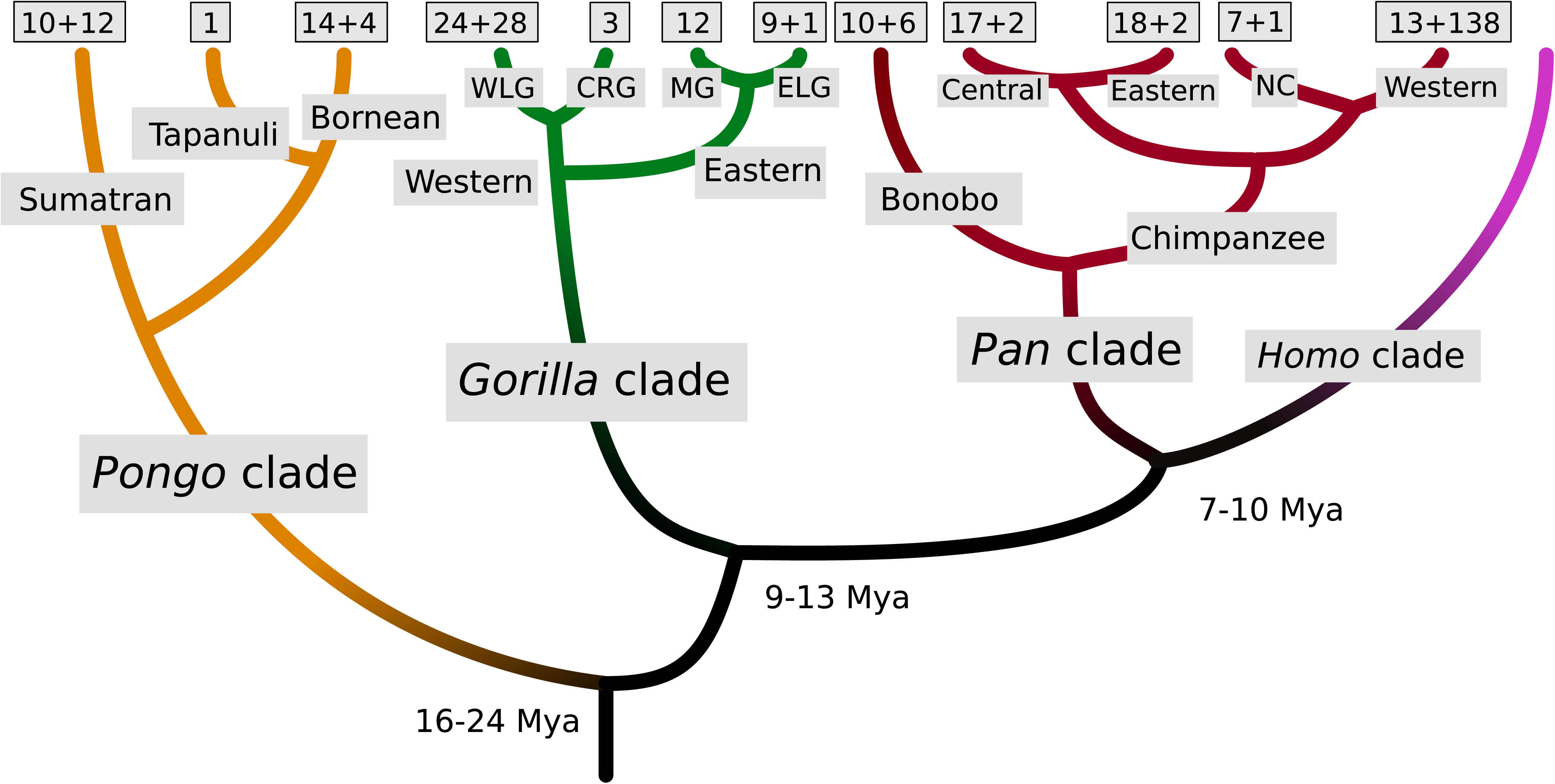
Phylogeny and overview of individuals from each of the clades. Orangutans (*Pongo*): Sumatran = *P. abelii*; Tapanuli = *P. tapanuliensis*; Bornean = *P. pygmaeus*. Gorillas (*Gorilla*): Western lowland gorilla (WLG) = *G. gorilla gorilla*; Cross-river gorilla (CRG) = *G. gorilla diehli*; Mountain gorilla (MG) = *G. beringei beringei*; Eastern lowland gorilla (ELG) = *G. beringei graueri*. Chimpanzees and bonobos (*Pan*): Bonobo = *P. paniscus*; Central chimpanzee = *P. troglodytes troglodytes*; Eastern chimpanzee = *P. troglodytes schweinfurthii*; Nigeria-Cameroon chimpanzee (NC) = *P. troglodytes ellioti*; Western chimpanzee = *P. troglodytes verus*. Divergence times in million years ago (Mya) from previous studies. Numbers of individuals as reference panel plus (+) captive panel.

In many studies of human and hominin evolution, great apes are used for comparison, either as outgroup when calculating statistics^30^ or for demographic modelling^31^, or with the purpose to put diversity or variation into context^5,32–36^. However, in many cases the great ape reference genomes are used for such purposes rather than diversity data, even though it is often important to put variation within species into context. For example, examining regions of homozygosity^37^ or inbreeding patterns in archaic hominins are better understood by variation data from great ape species^38^. Furthermore, information on great ape and primate variation guides studies on human disease^39^ and can be used to predict the effects of mutations in present-day people^40^.

Given the importance of such data for human-related research questions, we suggest that it is necessary to provide a comprehensive, curated panel of published great ape genomes. Importantly, this should contain information on non-variant sites within populations, to ascertain the status of each position in the genome. A considerable number of individuals is available as short-read sequencing data (*i.e.* using the Illumina platform), while long-read sequencing data was primarily used for reference genome assemblies. Hence, here we focus on the single-nucleotide variant (SNV) diversity, providing a coherent dataset for 332 high-coverage genomes from all extant great ape species^41^. Given that great apes are very closely related to humans and each other, all data was mapped to the human reference genome (GrCH38)^19^. We provide both the mappings (in CRAM format) and whole-genome variant call files (gVCF files), as well as sets of segregating sites across clades, all of which we hope will be a useful resource for numerous studies on human and primate genomics.

## Methods

### Samples

We used publicly available great ape genomes published in different studies (Table 1). These entail 23 chimpanzees, 12 bonobos, 27 gorillas and 10 orangutans from a landmark study on great ape diversity^18^; 32 chimpanzees^19^, 21 gorillas^20–22^ and 15 orangutans^23^ from subsequent population-scale studies on different wild-born individuals; and a captive panel of in total 143 chimpanzees, 4 bonobos, 29 gorillas and 16 orangutans from multiple studies with different focus^10,11,24–29,42–47^. All sequencing data was publicly available on the Sequencing Read Archive (SRA), and obtained through the European Nucleotide Archive branch for this study (Table S1). We did not consider individuals from studies reporting partial genomic data (such as chromosome 21^16^ or the exome^48^) or with insufficient sequencing coverage^17,49^ (below 12-fold, on average, across the genome). In some of the studies considered here, sequencing data was reported for additional individuals^18–20,23,46^, which we excluded due to low average coverage, or reported evidence of cross-contamination^18^. In the case of one individual (SAMEA104361539^23^), no data was available for one sequencing run accession (ERX2240355), leading to insufficient coverage. Finally, we merged data for identical individuals published using different identifiers or in different studies in order to reach sufficient coverage (see section “Captive panel” in Technical Validation). We only considered data generated through Illumina short-read sequencing, in an effort of building a coherent dataset. We note that both long-read and short-read sequencing data was generated for some of the individuals from which the most recent genome assemblies were generated – in such cases the short-read data are included here^10,11^.

**Table 1.** Overview of studies and numbers of individuals from each clade represented. Sequencing Read Archive (SRA) study identifiers and corresponding references are given in the first column. For metadata and per-sample identifiers see Tables S1-2.

|  | *Pan* |  | *Gorilla* |  | *Pongo* |  |  |
| --- | --- | --- | --- | --- | --- | --- | --- |
| Study accession | Chimpanzee | Bonobo | Western | Eastern | Sumatran | Bornean | Tapanuli |
| PRJNA189439<sup>18</sup> | 21 | 12 | 24 | 3 | 5 | 5 |  |
| PRJEB15086<sup>19</sup> | 32 |  |  |  |  |  |  |
| PRJEB19688<sup>23</sup> |  |  |  |  | 5 | 9 | 1 |
| PRJEB3220<sup>21</sup> |  |  |  | 12 |  |  |  |
| PRJEB12821<sup>22</sup> |  |  |  | 6 |  |  |  |
| PRJEB60463<sup>20</sup> |  |  | 3 |  |  |  |  |
| PRJNA563344<sup>28</sup> | 2 |  | 2 |  |  | 1 |  |
| PRJEB59576<sup>29</sup> | 4 |  | 19 | 1 | 1 | 1 |  |
| PRJEB29710<sup>26</sup> | 4 |  | 4 |  | 3 |  |  |
| PRJEB5937<sup>25</sup> | 9 |  |  |  |  |  |  |
| PRJDB3537<sup>27</sup> | 3 |  |  |  |  |  |  |
| PRJNA369439<sup>11</sup> |  |  |  |  | 1 |  |  |
| PRJNA526933<sup>10</sup> |  | 1 |  |  |  |  |  |
| PRJNA986878,<br>PRJNA986879,<br>PRJNA976699,<br>PRJNA976700,<br>PRJNA976701,<br>PRJNA602326<sup>24</sup> | 1 | 3 | 3 |  | 2 | 1 |  |
| PRJNA785018<sup>42</sup> |  |  |  |  |  | 1 |  |
| PRJEB1675<sup>45</sup> | 5 |  |  |  | 5 |  |  |
| PRJEB21589<sup>43</sup> | 1 |  |  |  |  |  |  |
| PRJNA856315<sup>46</sup> | 77 |  |  |  |  |  |  |
| PRJNA635393<sup>47</sup> | 39 |  |  |  |  |  |  |
| **Total** | **198** | **16** | **55** | **22** | **22** | **18** | **1** |

### Bioinformatic processing

Raw fastq files were downloaded using sratoolkit (https://hpc.nih.gov/apps/sratoolkit.html, version 3.0.6), and fastQC^50^ was applied for initial quality assessment. Adapter trimming was performed with trimmomatic (version 0.39)^51^. Reads were then mapped to the human genome version GRCh38 (GCA_000001405.15_GRCh38_no_alt_analysis_set from the UCSC genome browser) with bwa mem (version 0.7.16a)^52^, sorted with samtools (version 1.14)^53^ and unmapped reads removed. Read groups were assigned with picardtools (version 2.21.4) (http://broadinstitute.github.io/picard/), and duplicated reads were marked with GATK (version 4.1.4.0)^54^ MarkDuplicatesSpark. Finally, reads from all sequencing libraries for each individual were merged with samtools merge into a single CRAM file. These files are reported in the associated dataset. For two individuals, sequencing depth was more than 200-fold^27^, resulting in excessive spurious heterozygous calls. For coherence of the dataset, we restricted the analysis to a subset of the raw sequencing data (only one out of two run accessions each, as reported in Table S1).

Using this merged CRAM file per individual, genotypes were called per chromosome with GATK HaplotypeCaller, using the flag ‘-ERC GVCF’ to generate genome-wide VCF files. These files are reported in the associated dataset. For haploid sex chromosomes in male individuals, we performed haploid genotype calling.

For downstream analyses, we created joint callsets with GATK GenomicsDBImport, and GATK GenotypeGVCFs. After creating the callset for the wild-born individuals, we added the captive panels. We report sets of segregating sites per clade (*Pan*, *Gorilla*, *Pongo*) with and without the captive panel within the PHAIDRA repository^41^ (see Data Records section). Both sets are available as VCF files and in PLINK2 format after conversion using plink2^55^ (version 2.00a5) with the parameters ‘--max-alleles 2 --snps-only --make-pgen --maf 0.00’. A permissive set contains all information on segregating sites per individual. We also obtained a more stringent set after filtering using bcftools (version 1.21)^56^, retaining only bi-allelic SNPs passing a 36-basepair mappability filter^57^, excluding sites outside the central 98% of the coverage distribution per individual with a minimum of 5-fold coverage per site, and removing heterozygous positions with less than 15% of reads supporting one allele. We also report a joint set of segregating sites across all species^41^.

### Data analysis

We estimated depth of coverage using mosdepth^58^ (version 0.3.3) on the CRAM files, and used bcftools^56^ (version 1.16) to generate summary statistics on the VCF files. Genetic sexing was performed using ratios of the mean coverage per chromosome, with chrX:chr1 smaller than 0.75 and chrY:chrX larger than 0.1 to determine male sex.

We performed Principal Component Analysis (PCA) using VCF2PCACluster (version 1.41)^59^ on VCF files before and after filtering. For ADMIXTURE analyses, we subsampled 1,000,000 random autosomal loci from the VCF files and ran ADMIXTURE (version 1.3.0)^60^. Relatedness estimates were calculated using ngsRelate^61^ (version 2.0). Runs of Homozygosity were detected using bcftools roh^62^ (version 1.21) per chromosome per individual. Human contamination on captive individuals was estimated using HuConTest^63^. Subspecies assignment with f3-statistics was performed using admixtools2^64^. As outgroup, ancestral alleles were approximated by liftover of genomic coordinates to the macaque reference genome (rheMac10)^65^ using rtracklayer^66^ (version 1.58.0 in R version 4.2.2) and bedtools getfasta^67^ (version 2.31.1). Geolocalization of captive chimpanzees was performed using rareCAGA^16^ after liftover of genotypes to the human genome version hg19 with bcftools liftover^68^.

We used R^69^ versions 4.2.3 and 4.2.2 for plotting, with packages ggplot2^70^ (versions 3.4.4 and 3.5.1), gridExtra (version 2.3; 10.32614/CRAN.package.gridExtra), dplyr^71^ (version 1.1.4), tidyverse^72^ (version 2.0.0), ggh4x^73^ (version 0.2.8).

## Data Records

The full dataset is available through PHAIDRA with the University of Vienna, under the following DOI: https://doi.org/10.25365/phaidra.51441. This dataset contains the CRAM files (mapped reads) for 332 individuals, as well as gVCF files (genotype calls) for all 332 individuals for autosomes and X chromosomes, as well as Y chromosomes for the male individuals. Note that gVCF files are in the intermediate format provided by GATK HaplotypeCaller, which can be used for joint genotype calling but might be genotyped if used individually. For all files, md5sums are provided in Table S7. Furthermore, joint genotype calls in VCF format are available for the three species complexes *Pan*, *Gorilla* and *Pongo*, for the full set of genotype calls as well as a filtered set. A set of joint genotype call files across all 332 individuals are available on PHAIDRA^41^, as well as the EVA platform^74^ under the accession PRJEB97324^75^.

## Technical Validation

### Sequencing data

We report a curated dataset of previously published genomic data for 138 wild-born great ape individuals^18–23,25^, which constitute a reference panel for population genetic studies^41^. We also included 194 captive individuals from multiple studies^10,11,18,24–29,42,43,45–47^, resulting in a total dataset of 332 individuals. Only individuals with at least 12-fold average coverage across the genome were included, with a median of 23-fold and a maximum of 141-fold coverage (as obtained by mosdepth). Since in some cases the coverage of called genotypes was below this threshold, we also excluded three such individuals (SAMN02736775, SAMN01920524 and SAMEA104361528). Using the average coverage for the sex chromosomes, we report 208 (63%) of individuals as female, and 124 (37%) as male. We provide this information, alongside other summary metrics, in Table S2 and S3, as well as Fig. S1.

We obtained several quality control measures to ensure completeness of the data: average coverage per chromosome, the last position per chromosome, the numbers of non-reference records and heterozygous positions per chromosome, and the ratio of transitions to transversions per individual (Fig. S2-7). We present the genome-wide average coverage and heterozygosity in Fig. 2. Heterozygosity values recapitulate findings from previous studies when stratified by the different subspecies^3,18,19,21–23^. Furthermore, we estimated potential human contamination^63^ in captive individuals. We set a threshold of 1% in order to include individuals (Fig. 4A, Table S5), leading to exclusion of some individuals (SAMN29543728, SAMN29543727, SAMN29543724, SAMN29543729)^46^ with values above 1%.

**Figure 2.**
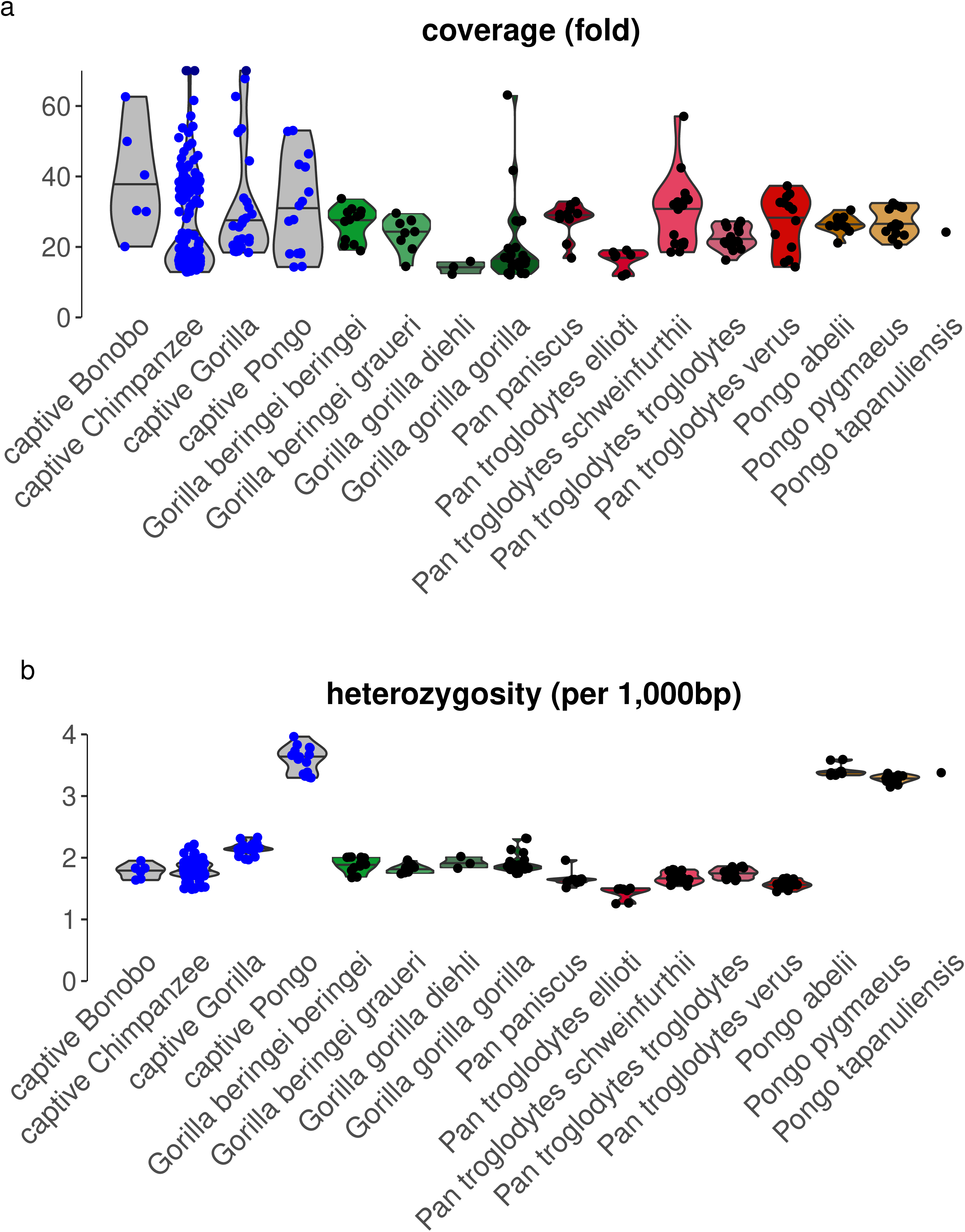
Distributions of coverage and heterozygosity. A) Average coverage per individual across subspecies. Note: captive *Pan* is cut at 70-fold for three samples with more than 100-fold coverage. B) Genome-wide heterozygosity per 1,000 base pairs (bp) per individual across subspecies.

Combined genotype calls of segregating sites per clade contain 117,472,161 sites for *Pan* (23,468,769 high quality sites after filtering), 80,907,619 sites for *Gorilla* (28,214,170 high quality sites after filtering), and 139,874,238 sites for *Pongo* (42,620,590 high quality sites after filtering). For downstream analyses, usually a coverage-based filtering is recommended. We report the central 98% of the coverage distribution for each individual, separately for autosomes and chromosome X in Table S4, with a lower cutoff of 5-fold coverage in cases where this value was below five. We advise the user to carefully consider additional filtering depending on their intended use of this dataset.

### Population genetic validation

We performed basic population genetic characterisation of the individuals in this dataset, which allows to assess the quality of the data in the context of previous findings. First, we performed a PCA, showing the expected population clustering of all individuals within the respective clades *Pan*, *Gorilla* and *Pongo* (Fig. 3A-C; Fig. S8-12). Captive individuals are shown in grey. Notably, a PCA on the unfiltered data shows outliers for the three orangutan individuals with the highest amount of human contamination in the sequencing data (see section below; Fig. S12). We also performed clustering with ADMIXTURE (Fig. 3D; Fig. S13-15), which recapitulates known patterns of subspecies stratification in these great ape species^3,18,19,21–23^.

**Figure 3.**
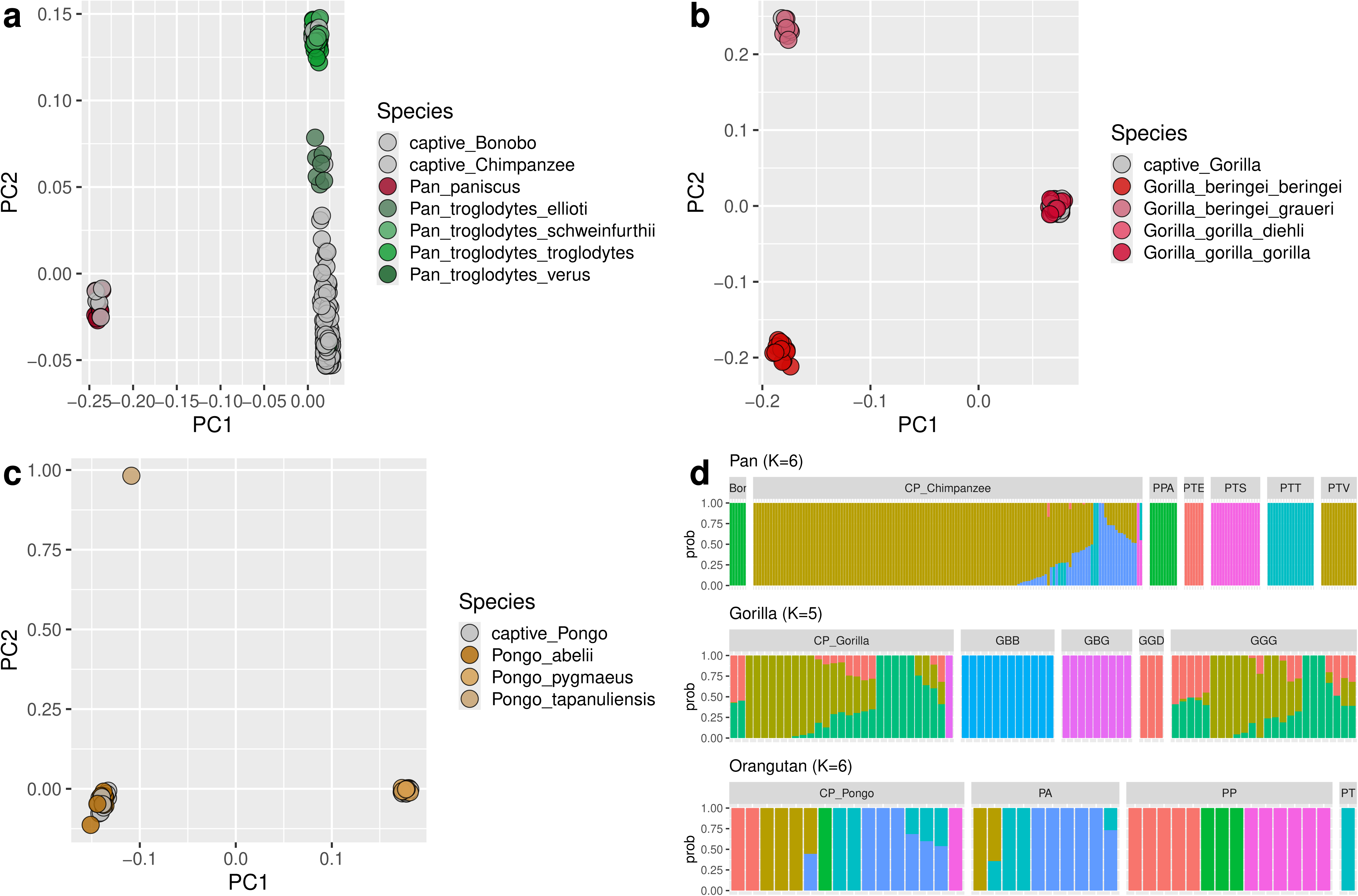
Basic population genetic characterisation of great ape individuals in this dataset. A) PCA for the *Pan* clade (chimpanzees and bonobos), B) PCA for the *Gorilla* clade, and C) PCA for the *Pongo* clade, all calculated on a filtered SNV callset. D) ADMIXTURE clustering for the *Pan*, *Gorilla* and *Pongo* clades (based on 1,000,000 random SNVs per clade). PPA, *Pan paniscus*; PTE, *Pan troglodytes ellioti*; PTS, *Pan troglodytes schweinfurthii*; PTT, *Pan troglodytes troglodytes*; PTV, *Pan troglodytes verus*; GBB, *Gorilla beringei beringei*; GBG, *Gorilla beringei graueri*; GGD, *Gorilla gorilla diehli*; GGG, *Gorilla gorilla gorilla*; PA, *Pongo abelii*; PP, *Pongo pygmaeus*; PT, *Pongo tapanuliensis*; CP, captive population.

Since we initially included all data reported in previous studies, we found several identical individuals based on relatedness estimates^61^ (KING relatedness larger than 0.4). No indication of identity was given in these respective publications. This affects the orangutan individual PD_0262/ORAN23^28,29^, as well as the gorilla individuals Banjo^18,26^, Mimi^18,26^, Mawenzi/PD_0264^26,29^ and PD_0189/PD_026^29^, each of which have two unique SRA biosample IDs. Furthermore, a total of 17 identical chimpanzee individuals sequenced in different studies were identified. Remarkably, the individual Donald^18^ appears to have been sequenced in three independent studies (as 4x0519^47^ and NS07602^46^). In most duplicate cases, the more recent study yielded high-coverage genomes (>30-fold), which we used for building this dataset. In some cases, in order to increase coverage we merged data after the additional step of inspecting heterozygosity. We report heatmaps of relatedness estimates including these identical individuals in the Supplementary Materials (Fig. S22-23), and provide a table of the biosample IDs for duplicated individuals (Table S5). We conclude that our data is comprehensive and reflects the original data published through these studies.

## Usage notes

Beyond the well-characterized datasets of wild-born great apes presented here as a reference dataset, we included 198 captive individuals from different studies. As described above, we assessed human contamination, retaining only individuals with less than 1% contamination. However, for three orangutan individuals, values close to 1% apparently still lead to false genotype calls and a shift in the PCA (Fig. S12). We conclude that quality filtering is recommended for subsequent analyses.

Furthermore, we provide an accurate assignment on the subspecies level based on f3-statistics (Table S6, Fig. S20)^76^, since 24 gorilla and 131 chimpanzee individuals did not have subspecies-level information in their SRA record, as well as two orangutans which were only labelled as *Pongo*. We assigned these two orangutans as *Pongo pygmaeus*, as reported in supplementary materials of a corresponding study^77^, though not in the SRA database. Most gorilla individuals are *Gorilla gorilla gorilla*, with the exception of PD_0179, a *Gorilla beringei graueri*, also reported as such only in the supplementary of the corresponding publication^29^ (Fig. 4B). Among chimpanzees, we identify PD_0259 and Rogger as *Pan troglodytes schweinfurthii*, CH114 as *Pan troglodytes troglodytes*, and 88A020 as *Pan troglodytes ellioti,* while all other chimpanzees are *Pan troglodytes verus.* The individual Donald/4x051946/NS0760245 is a known subspecies hybrid^18^. Furthermore, we performed geolocalisation of the captive chimpanzees (Fig. S21), finding, for example, an approximate origin of PD_0259 in northern Democratic Republic of Congo (Fig. 4C). We also identify 16 further individuals as likely subspecies hybrids in captivity (Fig. S21, Table S6). These analyses give a meaningful context for the genomes of these individuals, as they can complement diversity datasets of their respective subspecies or local population groups.

**Figure 4.**
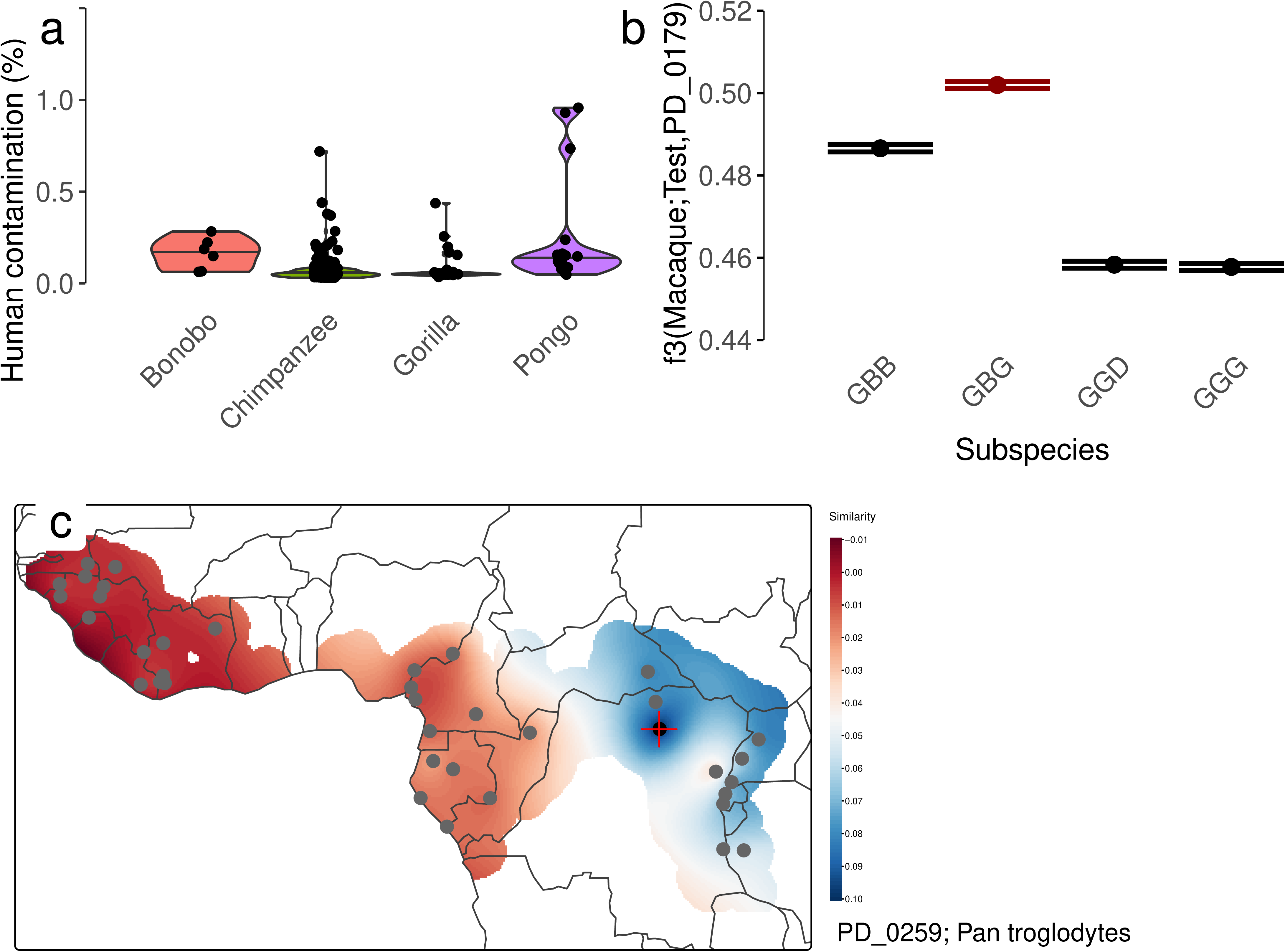
Characterizing the captive panel. A) Human contamination estimates in captive individuals. B) Subspecies assignment using f3-statistics for PD_0179. GBG, *Gorilla beringei beringei*; GBG, *Gorilla beringei graueri*; GGD, *Gorilla gorilla diehlii*; GGG, *Gorilla gorilla gorilla*. C) Geolocalization of a captive chimpanzee individual (PD_0259).

We also estimated pairwise relatedness between individuals^61^, recapitulating different degrees of background relatedness in some of the groups (Fig. S16-18), *e.g.* among bonobos (Fig. 5A) or Mountain gorillas (Fig. S17). Individuals from studies aimed at mutation rate estimates through trio sequencing^24–27^ were clearly identifiable by their first-degree relationships. Furthermore, multiple first-degree relationships were determined in captive chimpanzees. Known relationships are provided in Table S2, as well as inferred first-degree relationships (based on KING relatedness larger than 0.2), allowing to exclude such individuals from downstream analyses. Finally, we estimated runs of homozygosity^62^ (RoHs), a measure informative on long-term small effective population sizes, bottlenecks, and recent inbreeding^78^. We largely recapitulate previous findings^18^, *e.g.* more such RoHs in bonobos than chimpanzees or more in eastern gorillas than western lowland gorillas, while the captive individuals do not seem to show a systematic increase in RoHs (Fig. 5B; Fig. S19). Metadata are presented in Supplementary Tables and Figures, with Supplementary Figure legends in Table S8.

**Figure 5.**
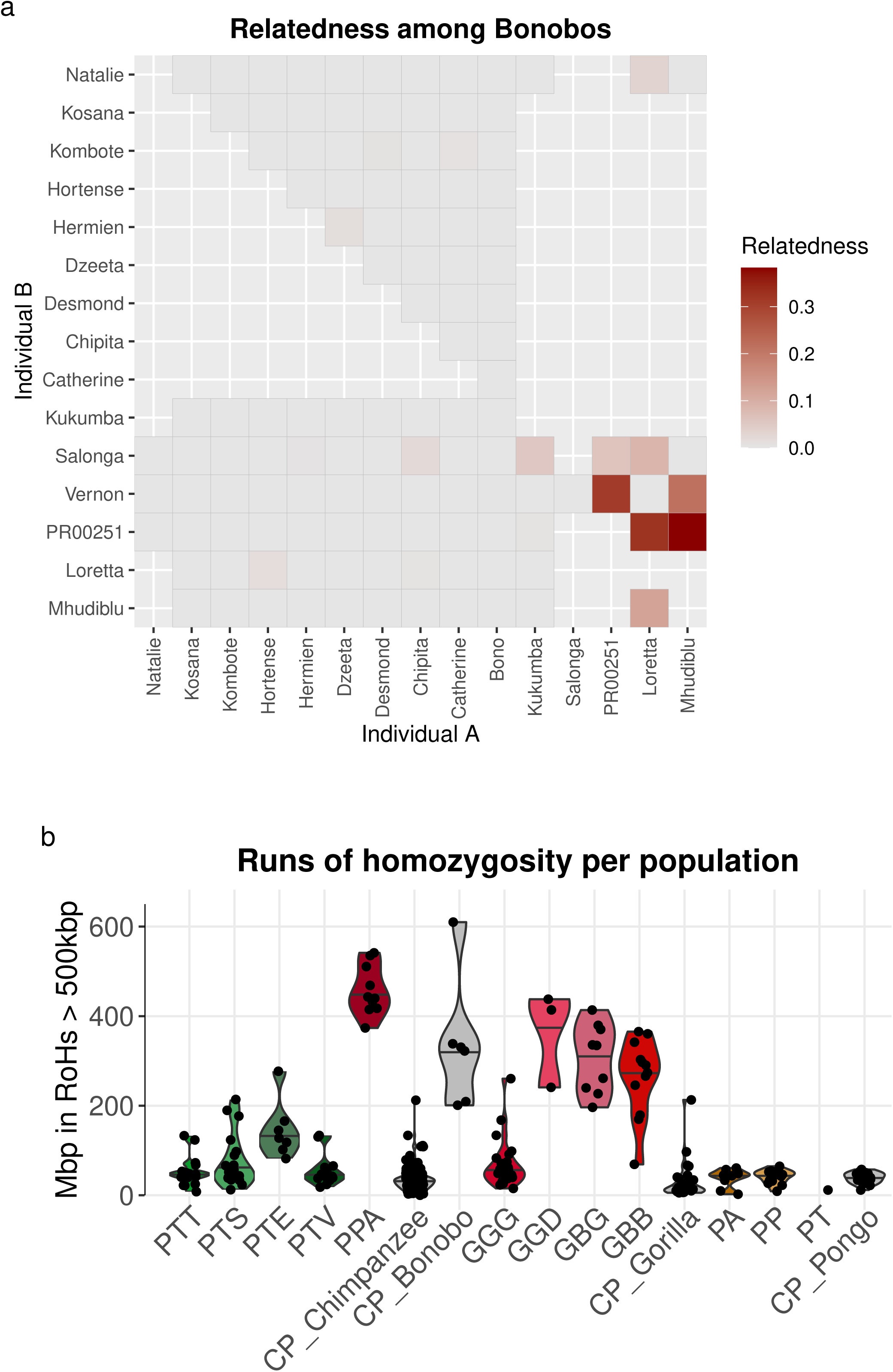
Kinship and runs of homozygosity. A) Relatedness among bonobos (*Pan paniscus*) as estimated by ngsRelate (KING method)^61^, as an example of individual relationships. Individuals from Trio sequencing (Mhudiblu, Loretta, PR00251) are distinguishable. Plots for all clades are found in Supplementary Materials. B) Runs of homozygosity as estimated by bcftools roh^62^ across all species and subspecies. For abbreviations see Table S2.

## Supplementary Tables

Table S1. Sequencing read archive (SRA) identifiers used to retrieve raw sequencing data for each individual.

Table S2. Metadata (species/subspecies, mean coverage, sex, relatedness, and other).

Table S3. Average coverage per chromosome.

Table S4. Central 98% coverage per individual.

Table S5. Reference table for duplicated individuals with their Biosample-IDs.

Table S6. Subspecies-assignment using f3-statistics, human contamination estimates for captive individuals, and geolocalization of chimpanzees.

Table S7. Md5sums for the CRAM and gVCF files in the dataset.

## Supplementary Figures

Figure S1. Coverage distribution per individual, stratified by subspecies.

Figure S2. Coverage per chromosome for 22 autosomes, stratified by subspecies.

Figure S3. Coverage for sex chromosomes, stratified by subspecies and sex (chrY for males only, mitochondrial genome (chrM) for all individuals).

Figure S4. Last called position per chromosome, ensuring completeness of data, stratified by subspecies.

Figure S5. Number of non-reference genotype calls per chromosome, stratified by subspecies.

Figure S6. Number of heterozygous positions per chromosome, stratified by subspecies.

Figure S7. Transition-to-transversion ratio per chromosome, stratified by individual.

Figure S8: Principal Component Analysis for 4 PCs all *Pan* individuals, calculated on unfiltered sites.

Figure S9: Principal Component Analysis for 4 PCs all *Pan troglodytes* (chimpanzee) individuals, calculated on unfiltered sites.

Figure S10: Principal Component Analysis for 4 PCs all *Pan paniscus* (bonobo) individuals, calculated on unfiltered sites.

Figure S11: Principal Component Analysis for 4 PCs all *Gorilla* individuals, calculated on unfiltered sites.

Figure S12: Principal Component Analysis for 4 PCs all *Pongo* individuals, calculated on unfiltered sites.

Figure S13: ADMIXTURE clustering for all *Pan* individuals, for 325 k=2 to k=10, calculated on 1,000,000 randomly chosen SNVs.

Figure S14: ADMIXTURE clustering for all *Gorilla* individuals, for k=1 to k=10, calculated on 1,000,000 randomly chosen SNVs.

Figure S15: ADMIXTURE clustering for all *Pongo* individuals, for k=1 to k=10, calculated on 1,000,000 randomly chosen SNVs.

Figure S16. Relatedness between all *Pan* individuals, as determined by KING in ngsRelate. Calculated on filtered dataset.

Figure S17. Relatedness between all *Gorilla* individuals, as determined by KING in ngsRelate. Calculated on filtered dataset.

Figure S18. Relatedness between all *Pongo* individuals, as determined by KING in ngsRelate. Calculated on filtered dataset.

Figure S19. Runs of Homozygosity (RoHs) in all individuals, as estimated by bcftools roh, stratified by subspecies and partitioned into short (50,000-250,000 bp), medium (250,000-1,000,000 bp) and long (>1,000,000) RoHs.

Figure S20. Subspecies assignment based on f3-statistics.

Figure S21. Geolocalization of chimpanzees based on rareCAGA.

Figure S22. Relatedness between all *Gorilla* individuals, including duplicated individuals that were removed from the dataset, as determined by KING in ngsRelate. Calculated on an unfiltered dataset.

Figure S23. Relatedness between all *Pongo* individuals, including duplicated individuals that were removed from the dataset, as determined by KING in ngsRelate. Calculated on an unfiltered dataset.

Figure S24. Relatedness between all *Pan* individuals, including duplicated individuals that were removed from the dataset, as determined by KING in ngsRelate. Calculated on an unfiltered dataset.

## Code Availability

The code used is available under https://github.com/admixVIE/Great_Ape_genomes.

## Data Availability

The dataset is available at https://doi.org/10.25365/phaidra.514, and has been deposited to EVA [PRJEB97324].

## Acknowledgements

We thank N. Schulmeister for revising parts of the data. This project has been funded by the Vienna Science and Technology Fund (WWTF) [10.47379/VRG20001] to M.K. S.H. was supported by the Austrian Science Fund (FWF) [10.55776/ESP546]. The computational results of this work have been achieved using supercomputer resources provided by the Vienna Scientific Cluster (VSC) and the Life Science Compute Cluster (LiSC) of the University of Vienna.

## Author contributions

M.K. conceived the study, analysed data, and wrote the manuscript with feedback from all coauthors. S.H., S.R. and X.H. analysed data.

