## Supplementary Figures for "A curated dataset of great ape genome diversity"

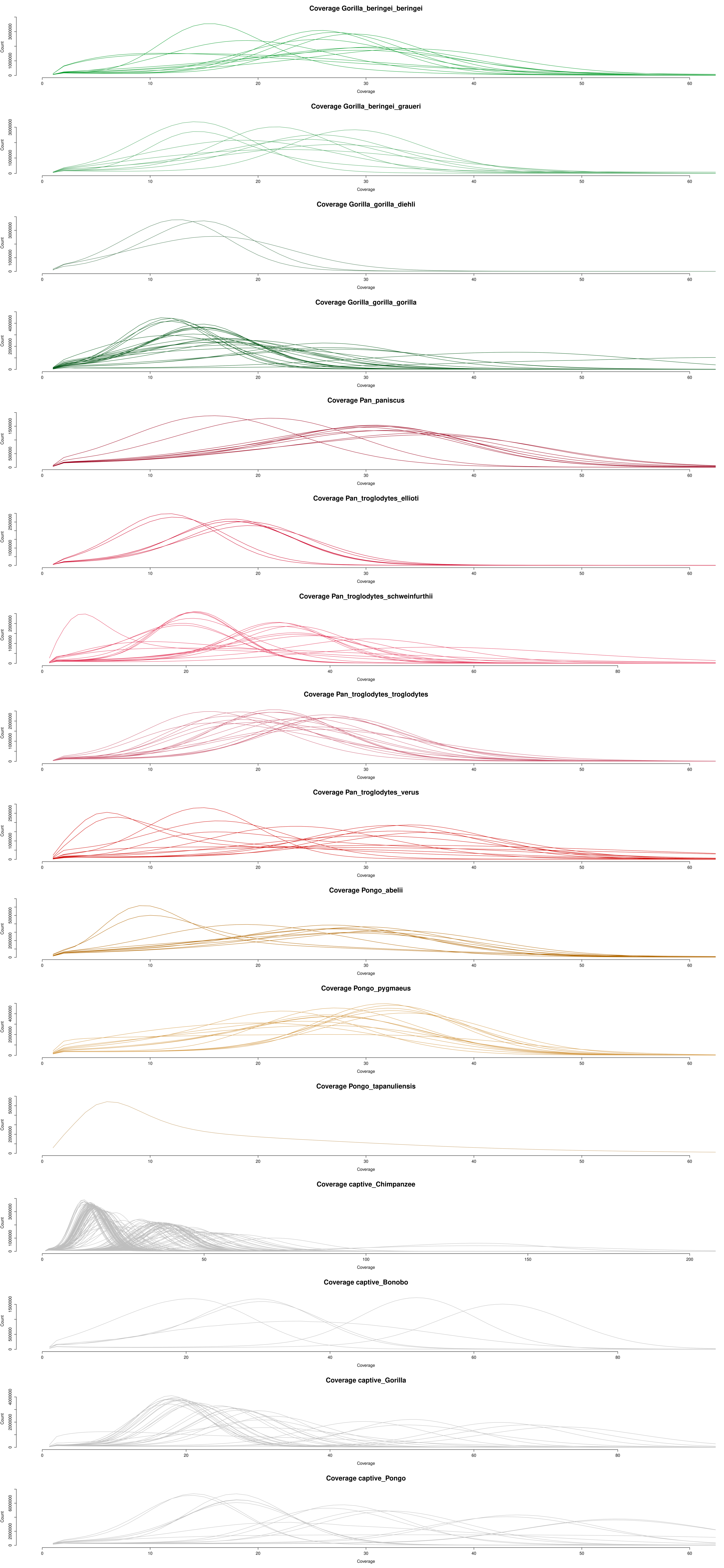
Coverage Gorilla_beringei_beringei
Coverage Gorilla_beringei_graueri
Coverage Gorilla_gorilla_diehli
Coverage Gorilla_gorilla_gorilla
Coverage Pan_paniscus
Coverage Pan_troglodytes_ellioti
Coverage Pan_troglodytes_schweinfurthii
Coverage Pan_troglodytes_troglodytes
Coverage Pan_troglodytes_verus
Coverage Pongo_abelii
Coverage Pongo_pygmaeus
Coverage Pongo_tapanuliensis
Coverage captive_Chimpanzee
Coverage captive_Bonobo
Coverage captive_Gorilla
Coverage captive_Pongo
Count
Coverage

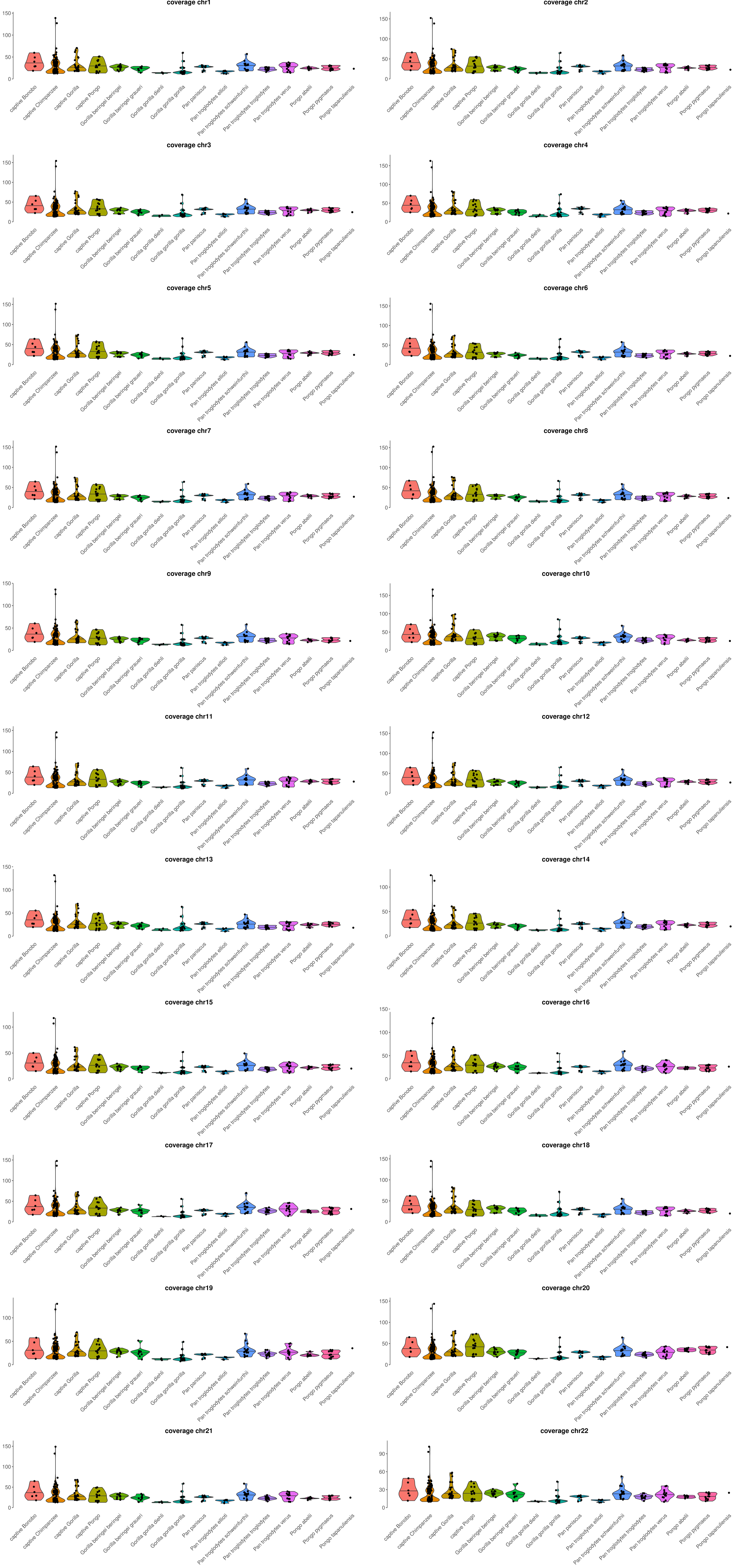
coverage chr1
coverage chr2
coverage chr3
coverage chr4
coverage chr5
coverage chr6
coverage chr7
coverage chr8
coverage chr9
coverage chr10
coverage chr11
coverage chr12
coverage chr13
coverage chr14
coverage chr15
coverage chr16
coverage chr17
coverage chr18
coverage chr19
coverage chr20
coverage chr21
coverage chr22
captive Bonobo
captive Chimpanzee
captive Gorilla
captive Pongo
Gorilla beringei beringei
Gorilla beringei graueri
Gorilla gorilla diehli
Gorilla gorilla gorilla
Pan paniscus
Pan troglodytes ellioti
Pan troglodytes schweinfurthii
Pan troglodytes troglodytes
Pan troglodytes verus
Pongo abelii
Pongo pygmaeus
Pongo tapanuliensis

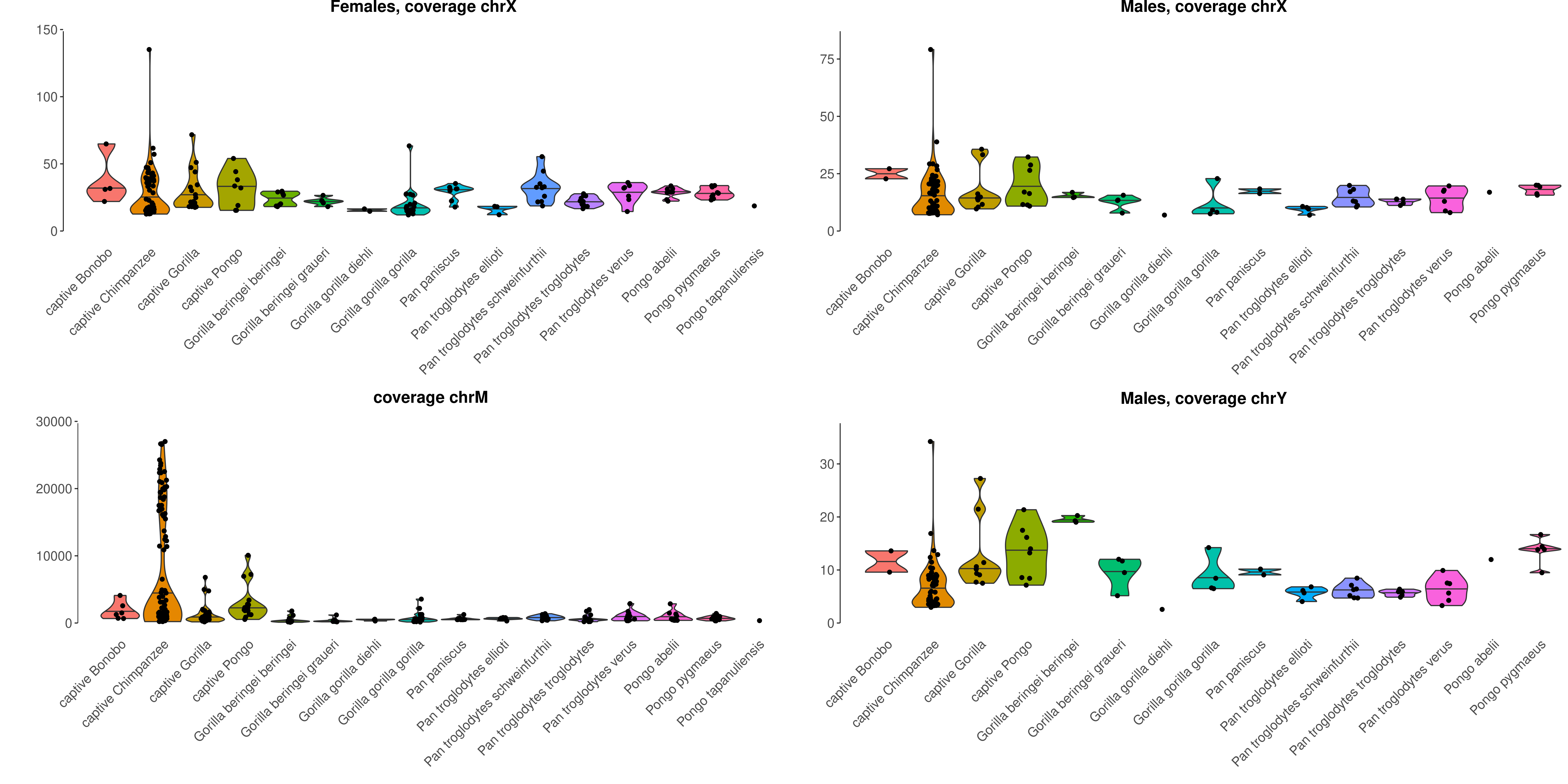

Females, coverage chrX
Males, coverage chrX
coverage chrM
Males, coverage chrY
150
100
50
0
75
25
30000
20000
10000
30
20
10
captive Bonobo
captive Chimpanzee
captive Gorilla
captive Pongo
Gorilla beringei beringei
Gorilla beringei graueri
Gorilla gorilla diehli
Gorilla gorilla gorilla
Pan paniscus
Pan troglodytes ellioti
Pan troglodytes schweinfurthii
Pan troglodytes troglodytes
Pan troglodytes verus
Pongo abelii
Pongo pygmaeus
Pongo tapanuliensis

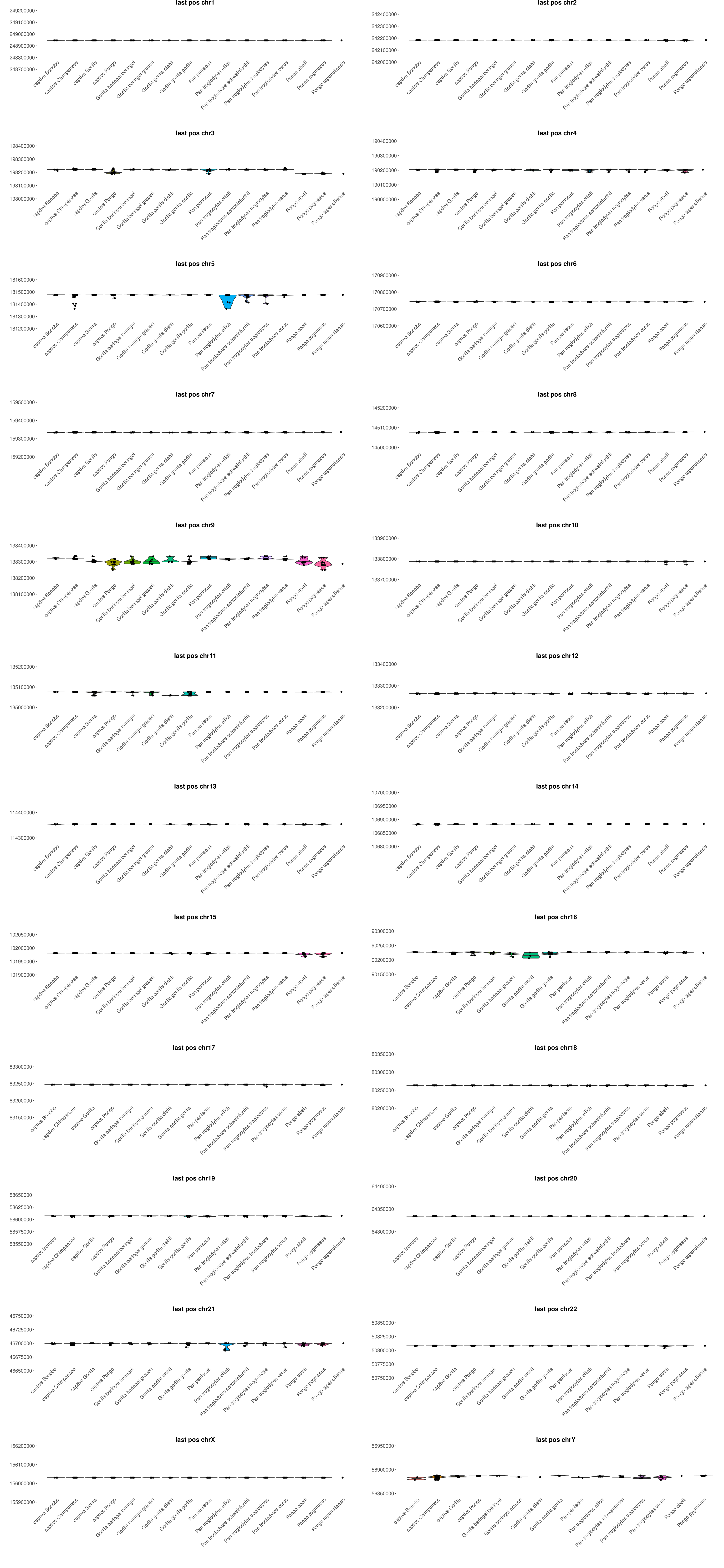
last pos chr1
249200000
249100000
249000000
248900000
248800000
248700000
last pos chr2
242400000
242300000
242200000
242100000
242000000
last pos chr3
198400000
198300000
198200000
198100000
198000000
last pos chr4
190400000
190300000
190200000
190100000
190000000
last pos chr5
181600000
181500000
181400000
181300000
181200000
last pos chr6
170900000
170800000
170700000
170600000
last pos chr7
159500000
159400000
159300000
159200000
last pos chr8
145200000
145100000
145000000
last pos chr9
138400000
138300000
138200000
138100000
last pos chr10
133900000
133800000
133700000
last pos chr11
135200000
135100000
135000000
last pos chr12
133400000
133300000
133200000
last pos chr13
114400000
114300000
last pos chr14
107000000
106950000
106900000
106850000
106800000
last pos chr15
102050000
102000000
101950000
101900000
last pos chr16
90300000
90250000
90200000
90150000
last pos chr17
83300000
83250000
83200000
83150000
last pos chr18
80350000
80300000
80250000
80200000
last pos chr19
58650000
58625000
58600000
58575000
58550000
last pos chr20
64400000
64350000
64300000
last pos chr21
46750000
46725000
46700000
46675000
46650000
last pos chr22
50850000
50825000
50800000
50775000
50750000
last pos chrX
156200000
156100000
156000000
155900000
last pos chrY
56950000
56900000
56850000
captive Bonobo
captive Chimpanzee
captive Gorilla
captive Pongo
Gorilla beringei beringei
Gorilla beringei graueri
Gorilla gorilla diehli
Gorilla gorilla gorilla
Pan paniscus
Pan troglodytes ellioti
Pan troglodytes schweinfurthii
Pan troglodytes troglodytes
Pan troglodytes verus
Pongo abelii
Pongo pygmaeus
Pongo tapanuliensis

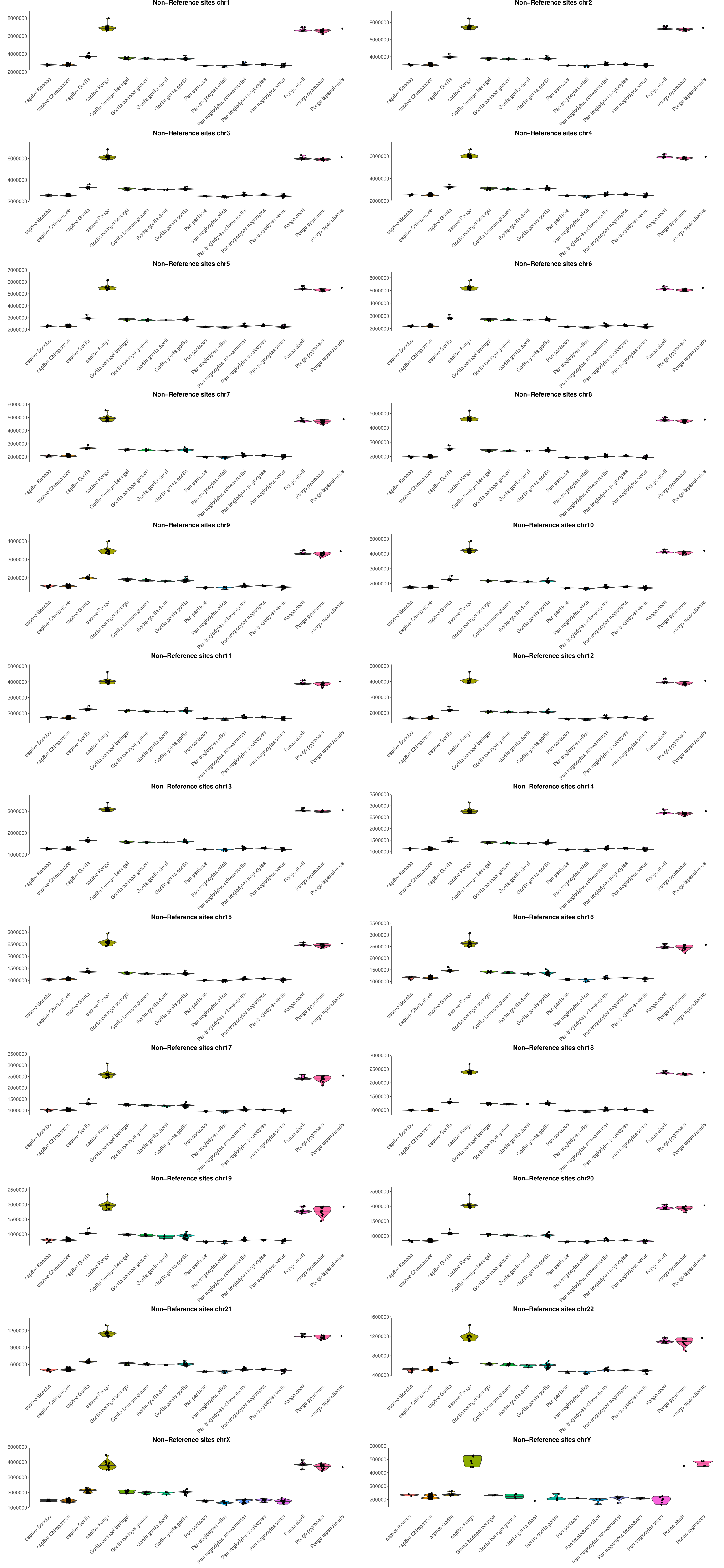

Non−Reference sites chr1
8000000
6000000
4000000
2000000
Non−Reference sites chr2
8000000
6000000
4000000
Non−Reference sites chr3
6000000
4000000
2000000
Non−Reference sites chr4
6000000
4000000
2000000
Non−Reference sites chr5
7000000
6000000
5000000
4000000
3000000
2000000
Non−Reference sites chr6
6000000
5000000
4000000
3000000
2000000
Non−Reference sites chr7
6000000
5000000
4000000
3000000
2000000
Non−Reference sites chr8
5000000
4000000
3000000
2000000
Non−Reference sites chr9
4000000
3000000
2000000
Non−Reference sites chr10
5000000
4000000
3000000
2000000
Non−Reference sites chr11
5000000
4000000
3000000
2000000
Non−Reference sites chr12
5000000
4000000
3000000
2000000
Non−Reference sites chr13
3000000
2000000
1000000
Non−Reference sites chr14
3500000
3000000
2500000
2000000
1500000
1000000
Non−Reference sites chr15
3000000
2500000
2000000
1500000
1000000
Non−Reference sites chr16
3500000
3000000
2500000
2000000
1500000
1000000
Non−Reference sites chr17
3500000
3000000
2500000
2000000
1500000
1000000
Non−Reference sites chr18
3000000
2500000
2000000
1500000
1000000
Non−Reference sites chr19
2500000
2000000
1500000
1000000
Non−Reference sites chr20
2500000
2000000
1500000
1000000
Non−Reference sites chr21
1200000
900000
600000
Non−Reference sites chr22
1600000
1200000
800000
400000
Non−Reference sites chrX
5000000
4000000
3000000
2000000
1000000
Non−Reference sites chrY
600000
500000
400000
300000
200000
captive Bonobo
captive Chimpanzee
captive Gorilla
captive Pongo
Gorilla beringei beringei
Gorilla beringei graueri
Gorilla gorilla diehli
Gorilla gorilla gorilla
Pan paniscus
Pan troglodytes ellioti
Pan troglodytes schweinfurthii
Pan troglodytes troglodytes
Pan troglodytes verus
Pongo abelii
Pongo pygmaeus
Pongo tapanuliensis

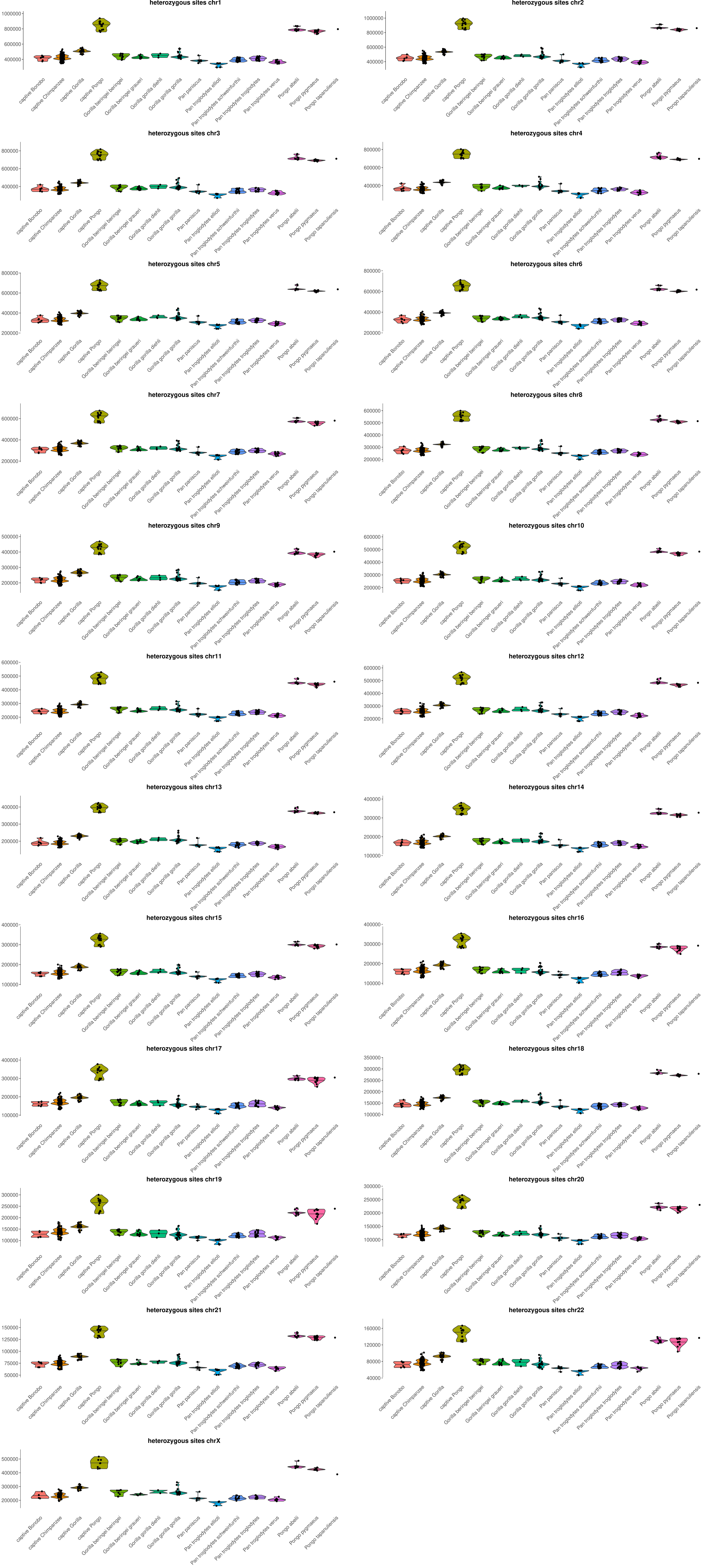
heterozygous sites chr1
heterozygous sites chr2
heterozygous sites chr3
heterozygous sites chr4
heterozygous sites chr5
heterozygous sites chr6
heterozygous sites chr7
heterozygous sites chr8
heterozygous sites chr9
heterozygous sites chr10
heterozygous sites chr11
heterozygous sites chr12
heterozygous sites chr13
heterozygous sites chr14
heterozygous sites chr15
heterozygous sites chr16
heterozygous sites chr17
heterozygous sites chr18
heterozygous sites chr19
heterozygous sites chr20
heterozygous sites chr21
heterozygous sites chr22
heterozygous sites chrX
captive Bonobo
captive Chimpanzee
captive Gorilla
captive Pongo
Gorilla beringei beringei
Gorilla beringei graueri
Gorilla gorilla diehli
Gorilla gorilla gorilla
Pan paniscus
Pan troglodytes ellioti
Pan troglodytes schweinfurthii
Pan troglodytes troglodytes
Pan troglodytes verus
Pongo abelii
Pongo pygmaeus
Pongo tapanuliensis

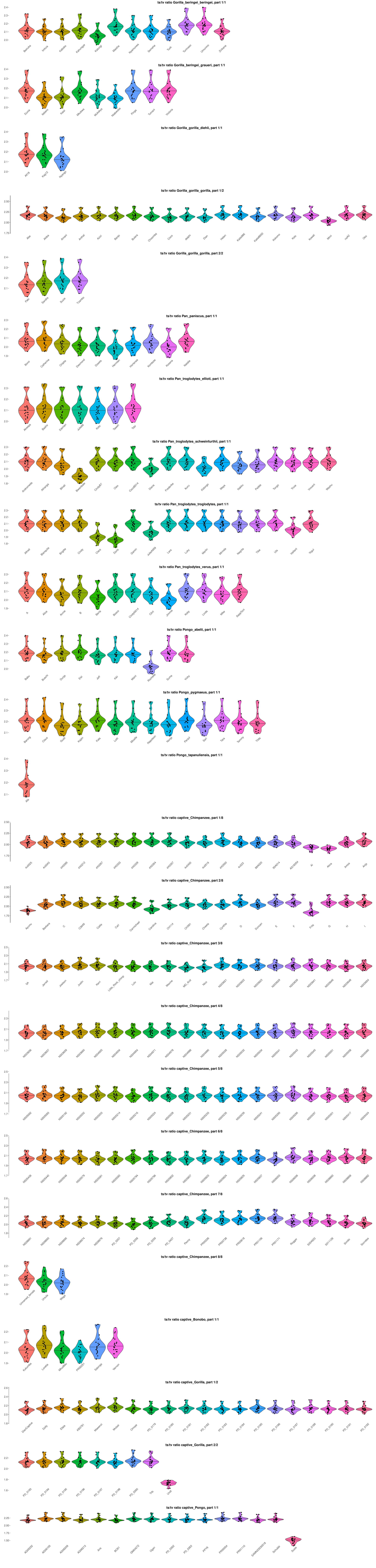

ts/tv ratio Gorilla_beringei_beringei, part 1/1
ts/tv ratio Gorilla_beringei_graueri, part 1/1
ts/tv ratio Gorilla_gorilla_diehli, part 1/1
ts/tv ratio Gorilla_gorilla_gorilla, part 1/2
ts/tv ratio Gorilla_gorilla_gorilla, part 2/2
ts/tv ratio Pan_paniscus, part 1/1
ts/tv ratio Pan_troglodytes_ellioti, part 1/1
ts/tv ratio Pan_troglodytes_schweinfurthii, part 1/1
ts/tv ratio Pan_troglodytes_troglodytes, part 1/1
ts/tv ratio Pan_troglodytes_verus, part 1/1
ts/tv ratio Pongo_abelii, part 1/1
ts/tv ratio Pongo_pygmaeus, part 1/1
ts/tv ratio Pongo_tapanuliensis, part 1/1
ts/tv ratio captive_Chimpanzee, part 1/8
ts/tv ratio captive_Chimpanzee, part 2/8
ts/tv ratio captive_Chimpanzee, part 3/8
ts/tv ratio captive_Chimpanzee, part 4/8
ts/tv ratio captive_Chimpanzee, part 5/8
ts/tv ratio captive_Chimpanzee, part 6/8
ts/tv ratio captive_Chimpanzee, part 7/8
ts/tv ratio captive_Chimpanzee, part 8/8
ts/tv ratio captive_Bonobo, part 1/1
ts/tv ratio captive_Gorilla, part 1/2
ts/tv ratio captive_Gorilla, part 2/2
ts/tv ratio captive_Pongo, part 1/1

### Pan PCA

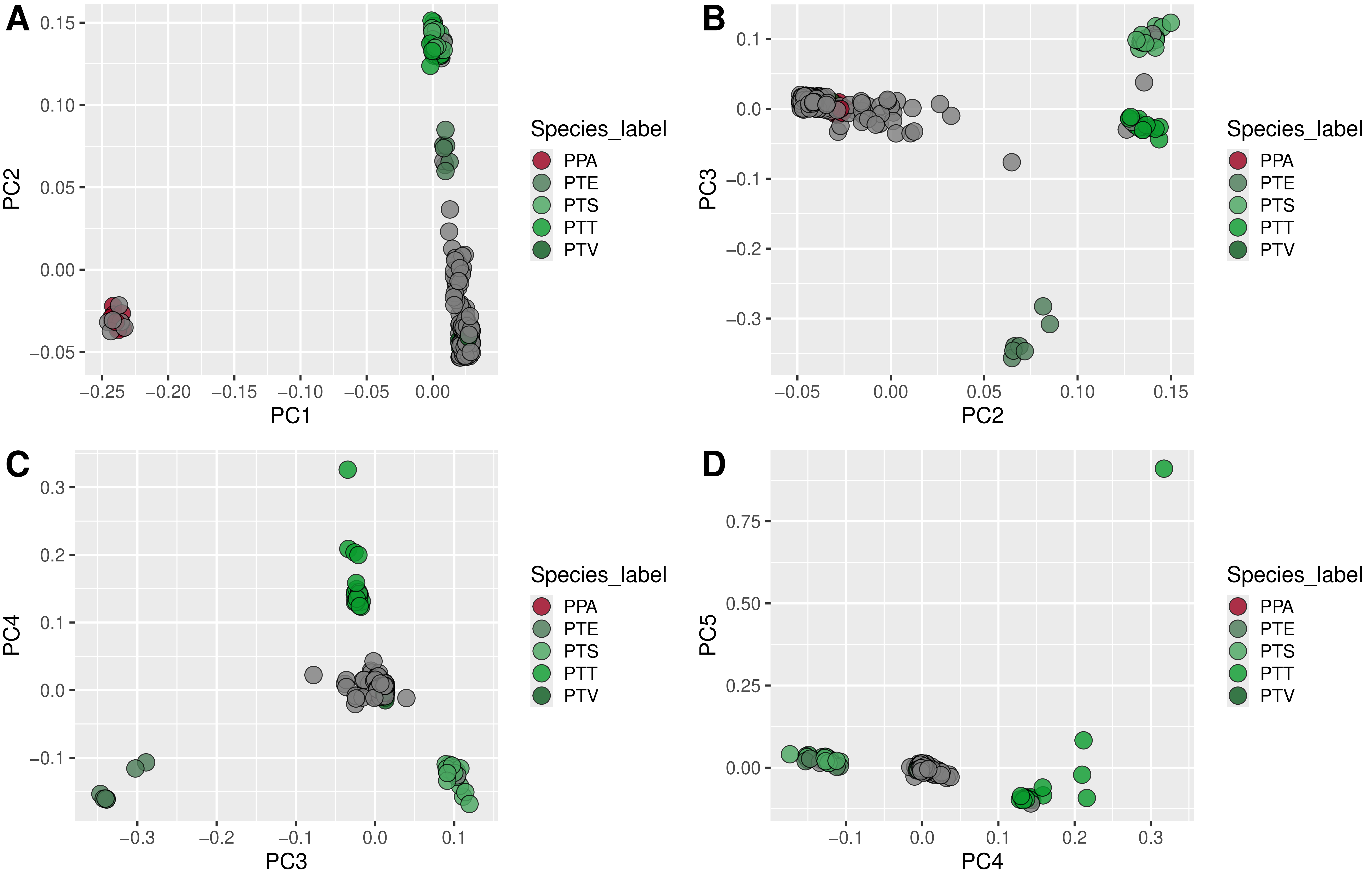

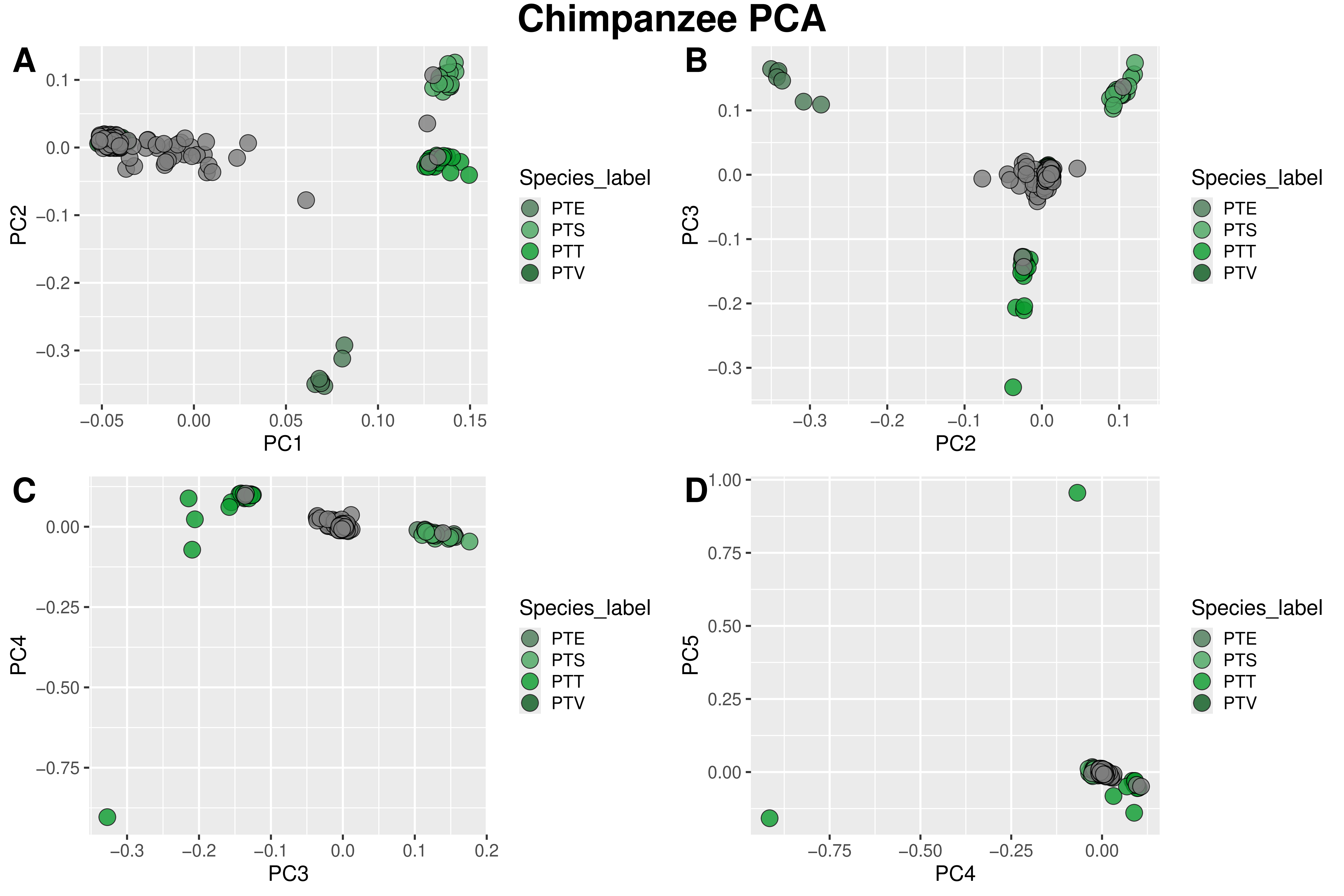

Chimpanzee PCA
A
B
C
D
PC1
PC2
PC3
PC4
PC5
Species_label
PTE
PTS
PTT
PTV

### Bonobo PCA

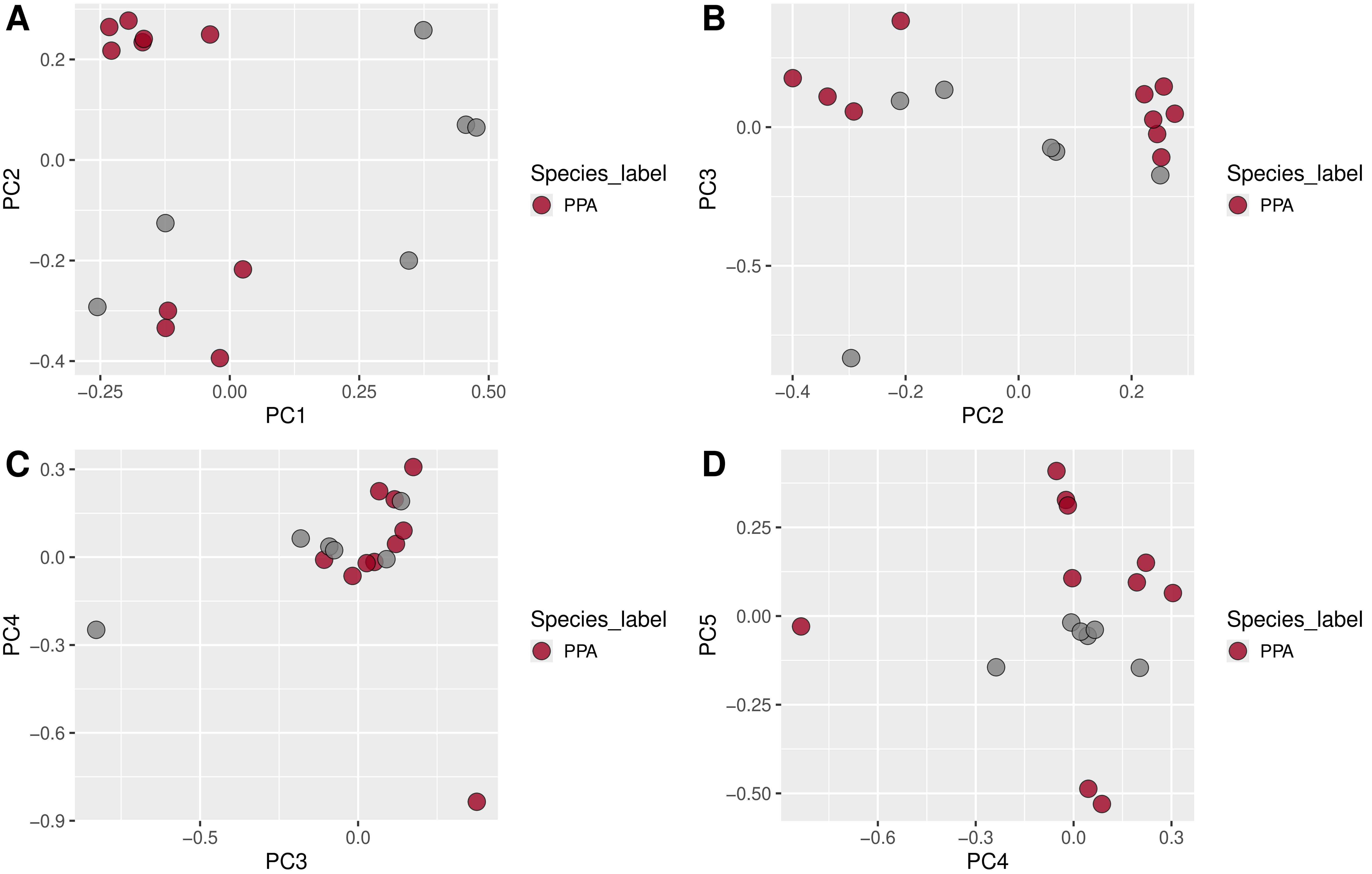
A
B
C
D
PC1
PC2
PC3
PC4
PC5
Species_label
PPA
Species_label
PPA
Species_label
PPA
Species_label
PPA
−0.25
0.00
0.25
0.50
0.2
0.0
−0.2
−0.4
−0.4
−0.2
0.0
0.2
0.0
−0.5
0.3
0.0
−0.3
−0.6
−0.9
−0.5
0.0
0.25
0.00
−0.25
−0.50
−0.6
−0.3
0.0
0.3

### Gorilla PCA

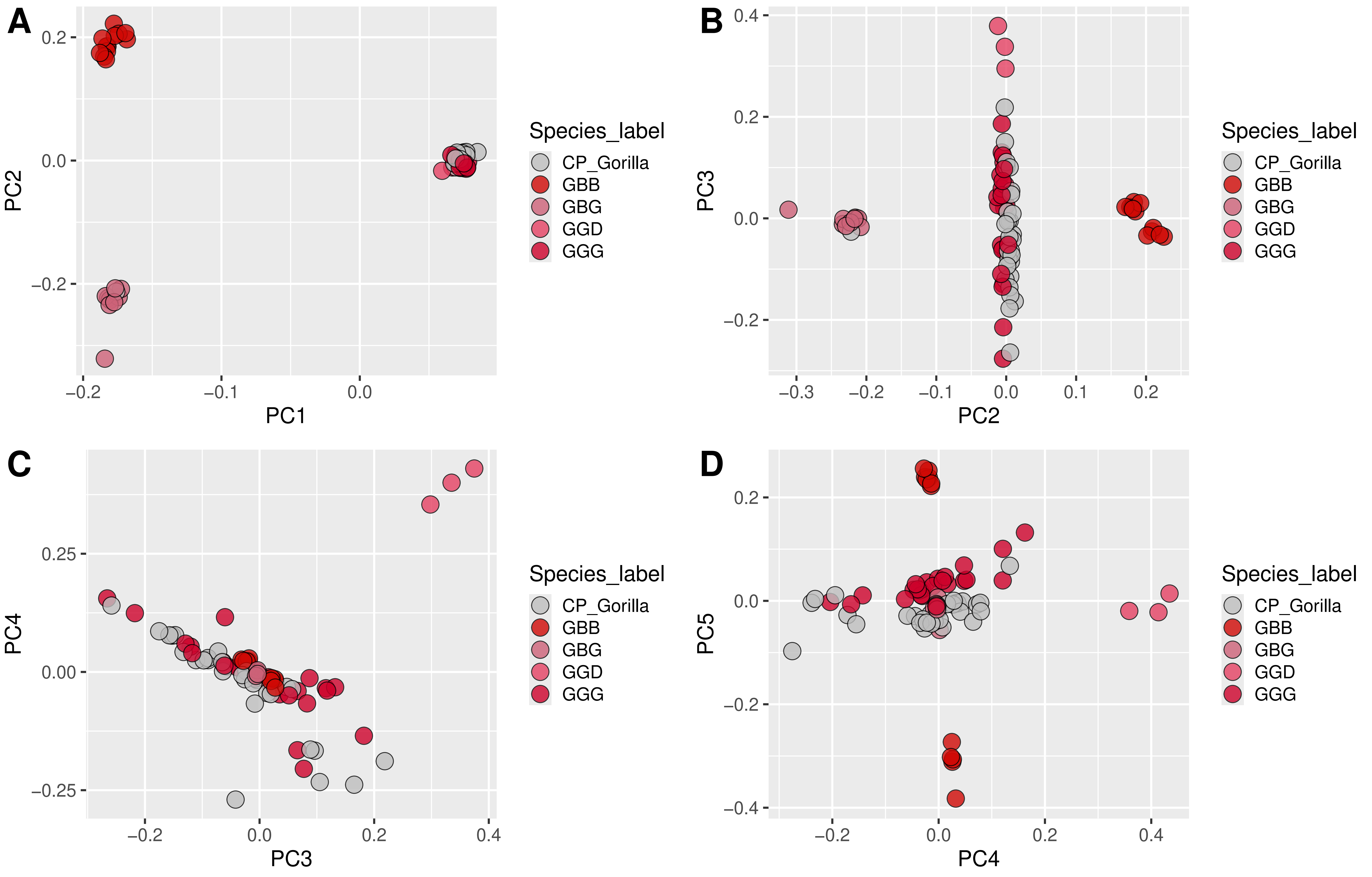

A
B
C
D
PC1
PC2
PC3
PC4
PC5
Species_label
CP_Gorilla
GBB
GBG
GGD
GGG

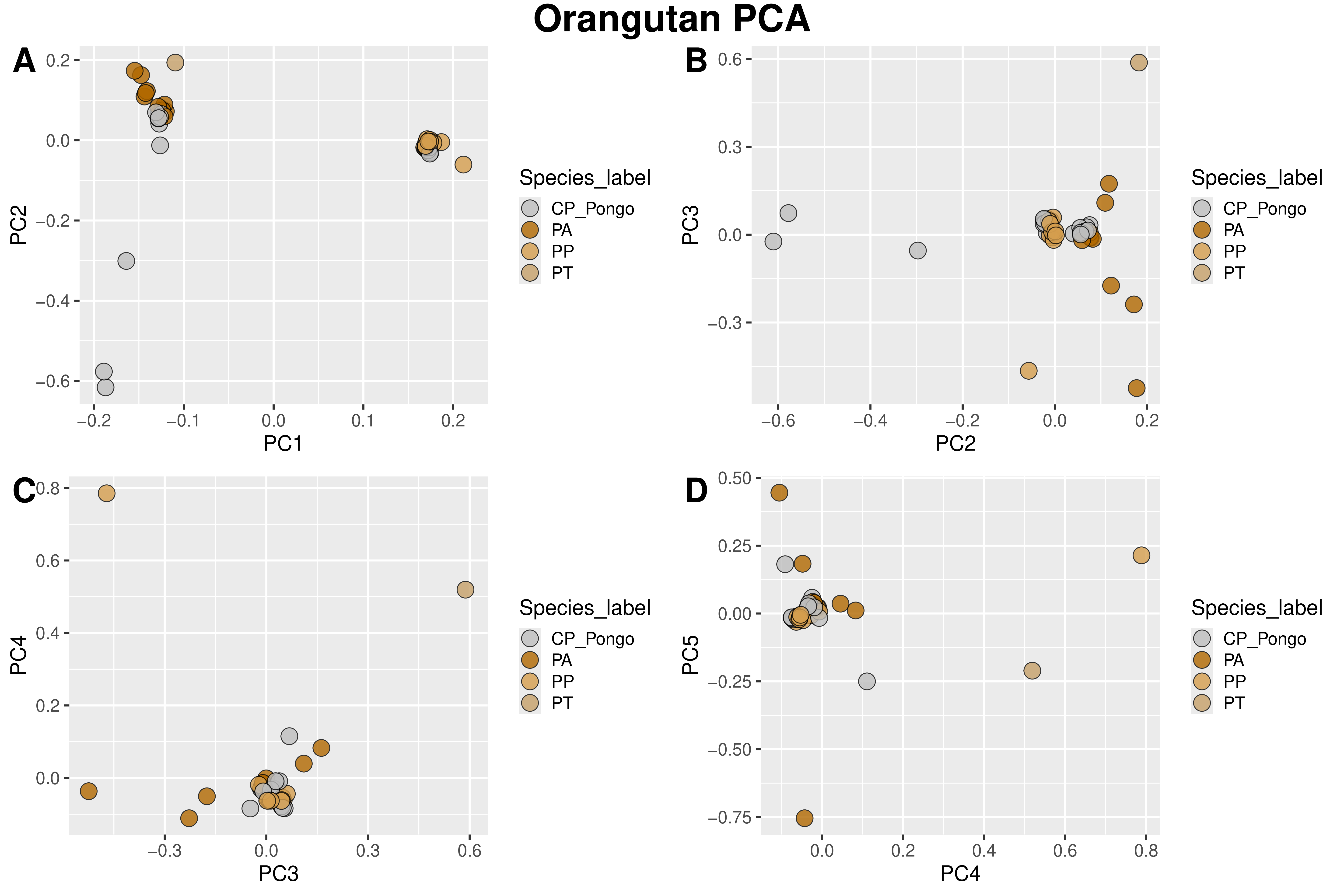
Orangutan PCA
A
B
C
D
PC1
PC2
PC3
PC4
PC5
Species_label
CP_Pongo
PA
PP
PT

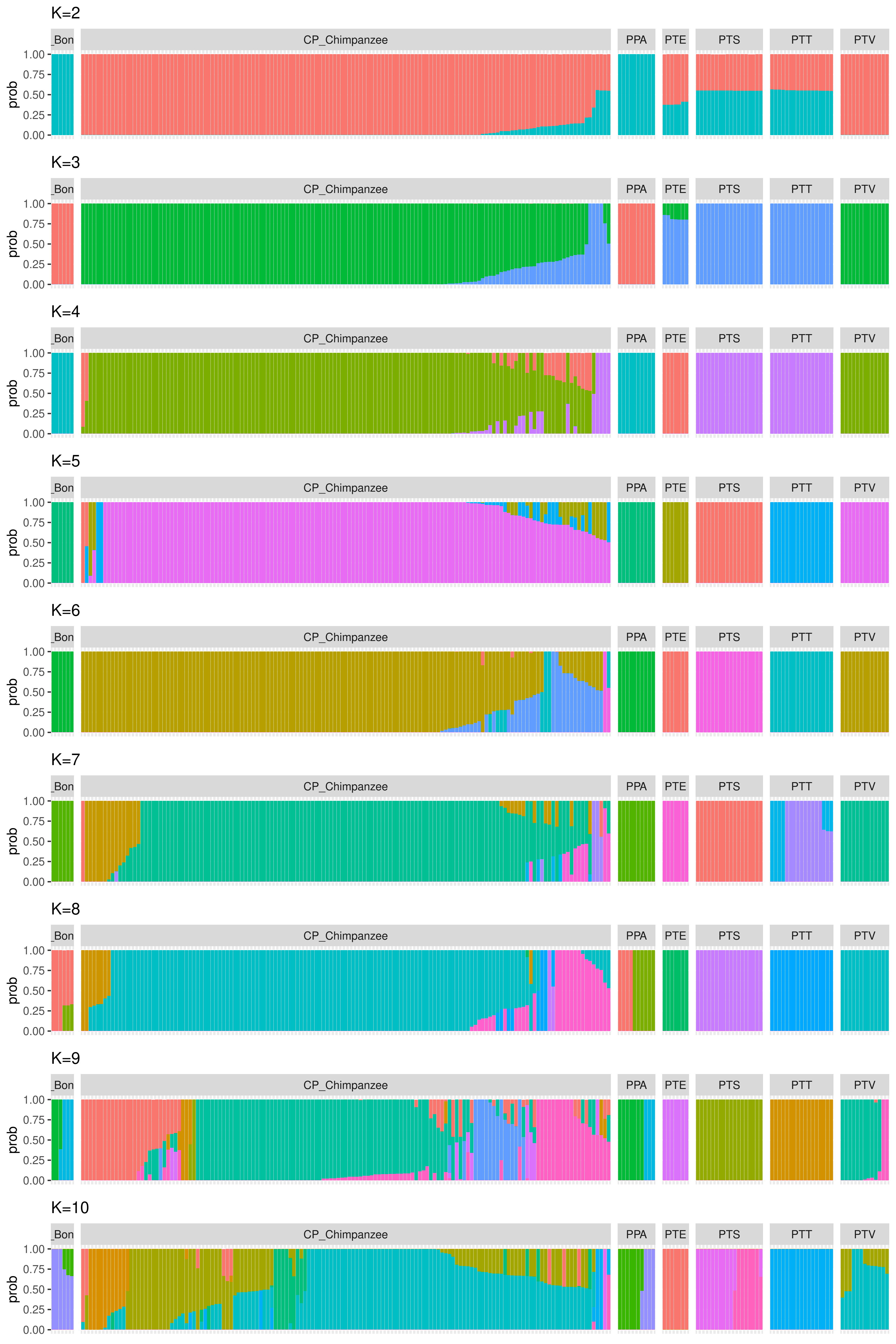
K=2
K=3
K=4
K=5
K=6
K=7
K=8
K=9
K=10
_Bon
CP_Chimpanzee
PPA
PTE
PTS
PTT
PTV
prob
1.00
0.75
0.50
0.25
0.00

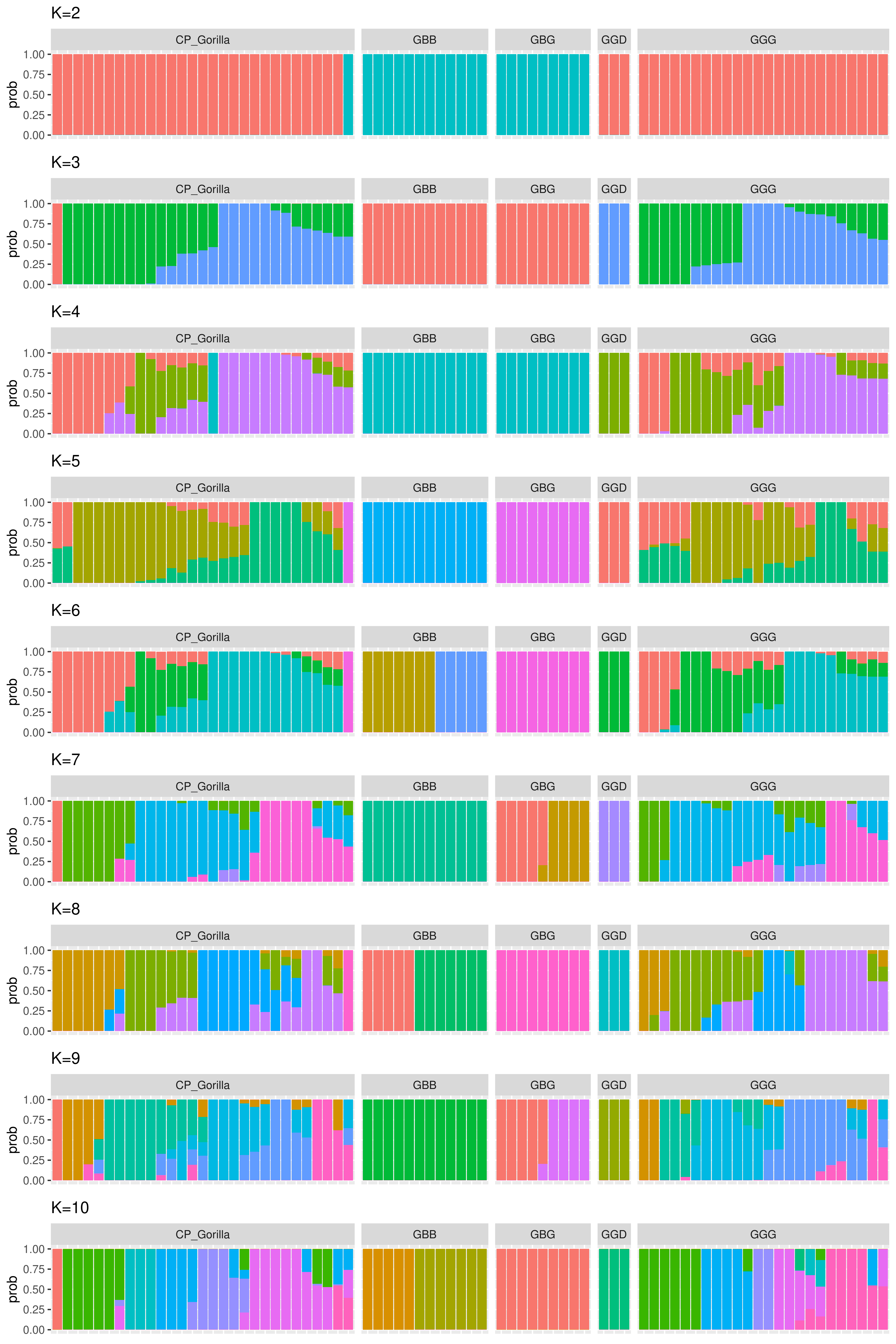
K=2
K=3
K=4
K=5
K=6
K=7
K=8
K=9
K=10
CP_Gorilla
GBB
GBG
GGD
GGG
prob
1.00
0.75
0.50
0.25
0.00

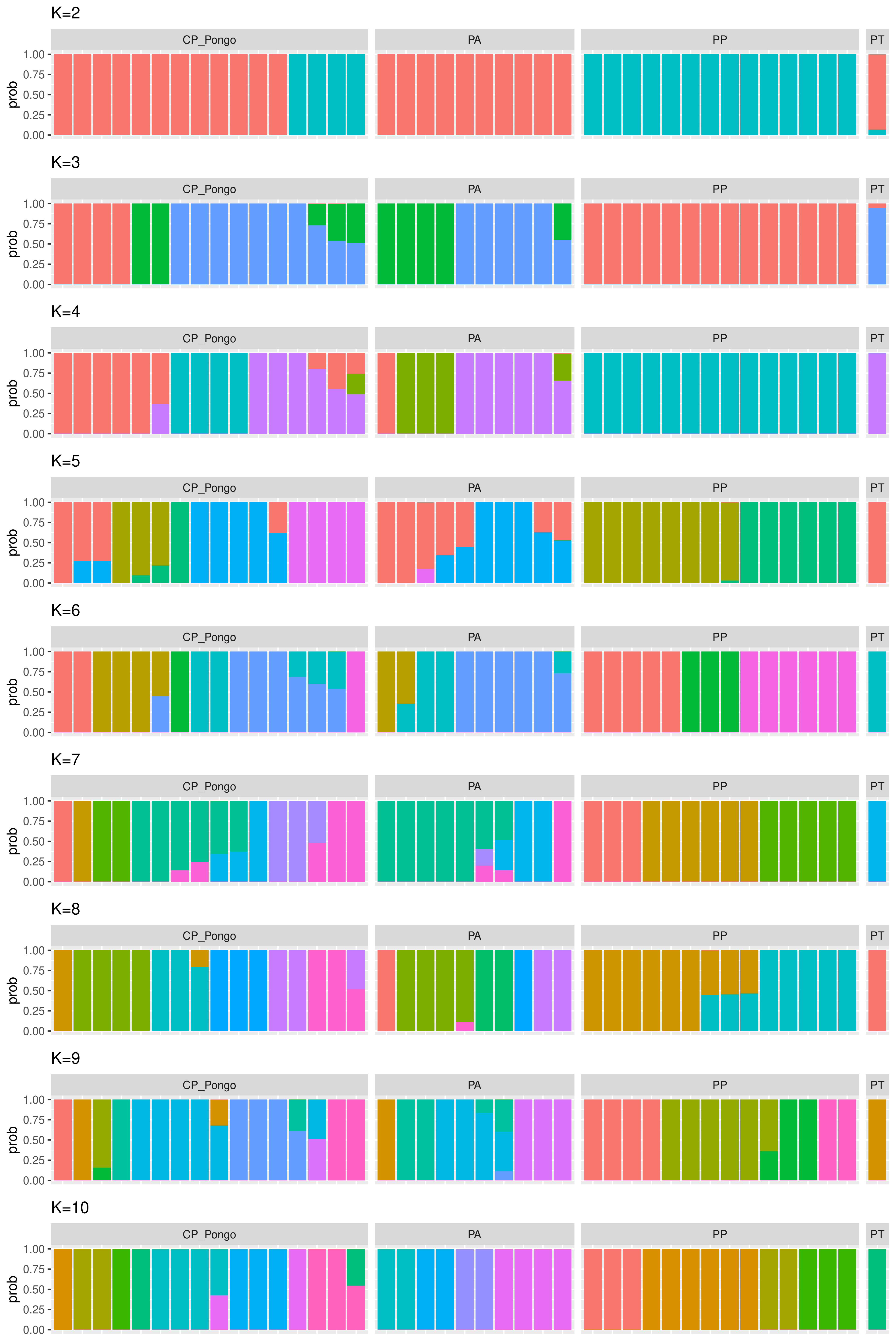

K=2
K=3
K=4
K=5
K=6
K=7
K=8
K=9
K=10
CP_Pongo
PA
PP
PT
prob
1.00
0.75
0.50
0.25
0.00

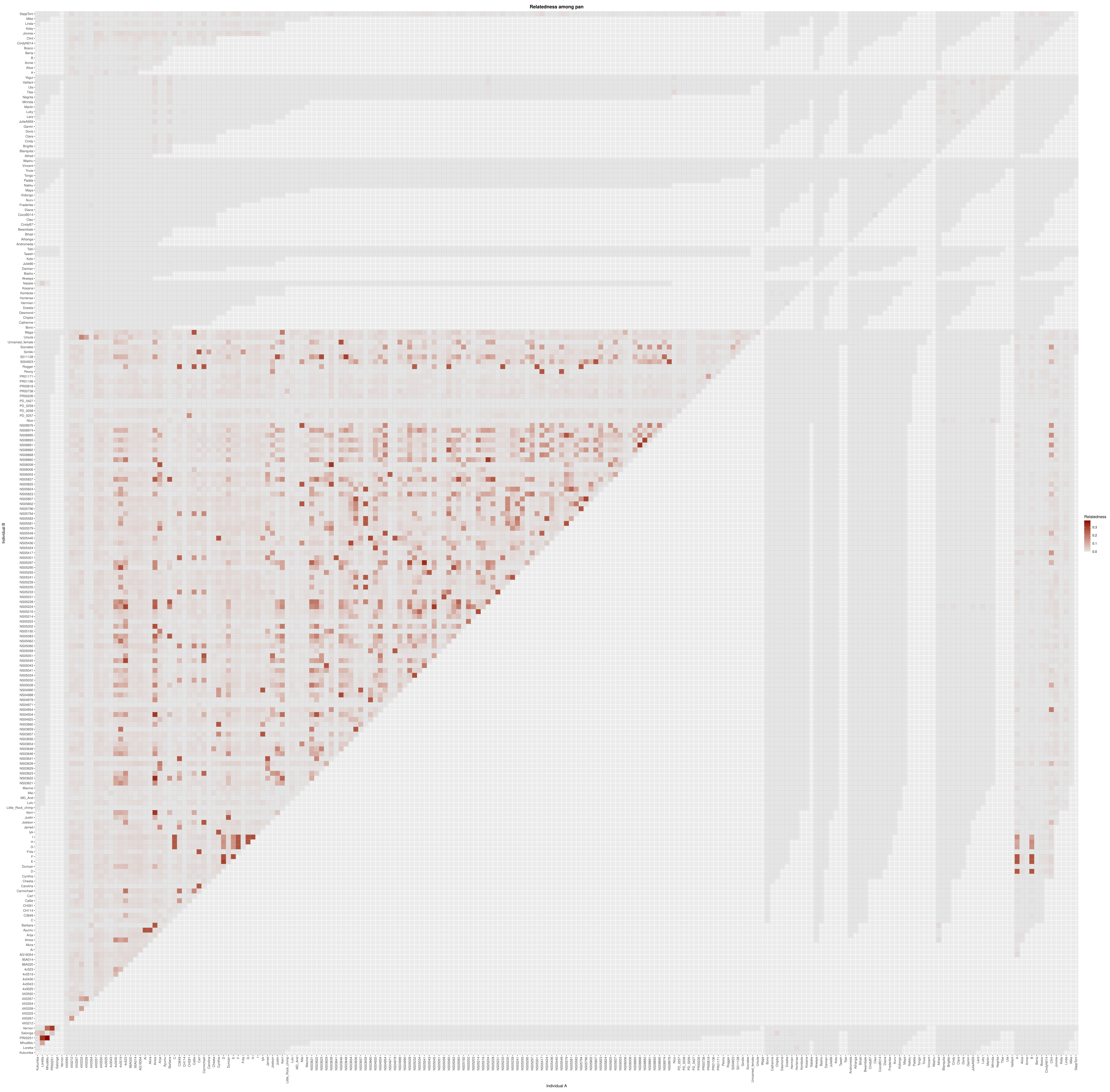
Relatedness among pan
Individual B
Individual A
Relatedness
0.3
0.2
0.1
0.0

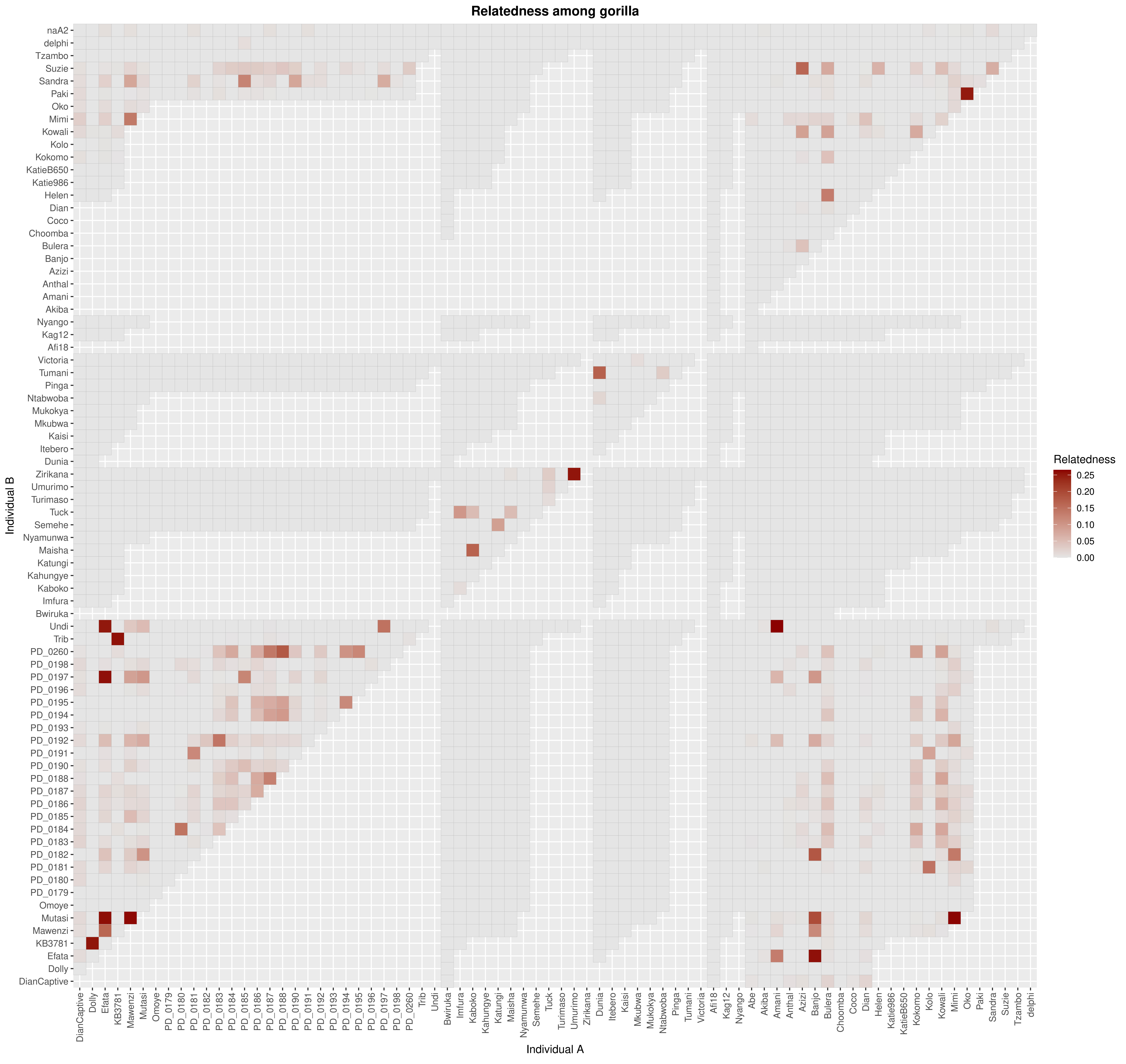

Relatedness among gorilla
Individual A
Individual B
Relatedness
0.25
0.20
0.15
0.10
0.05
0.00

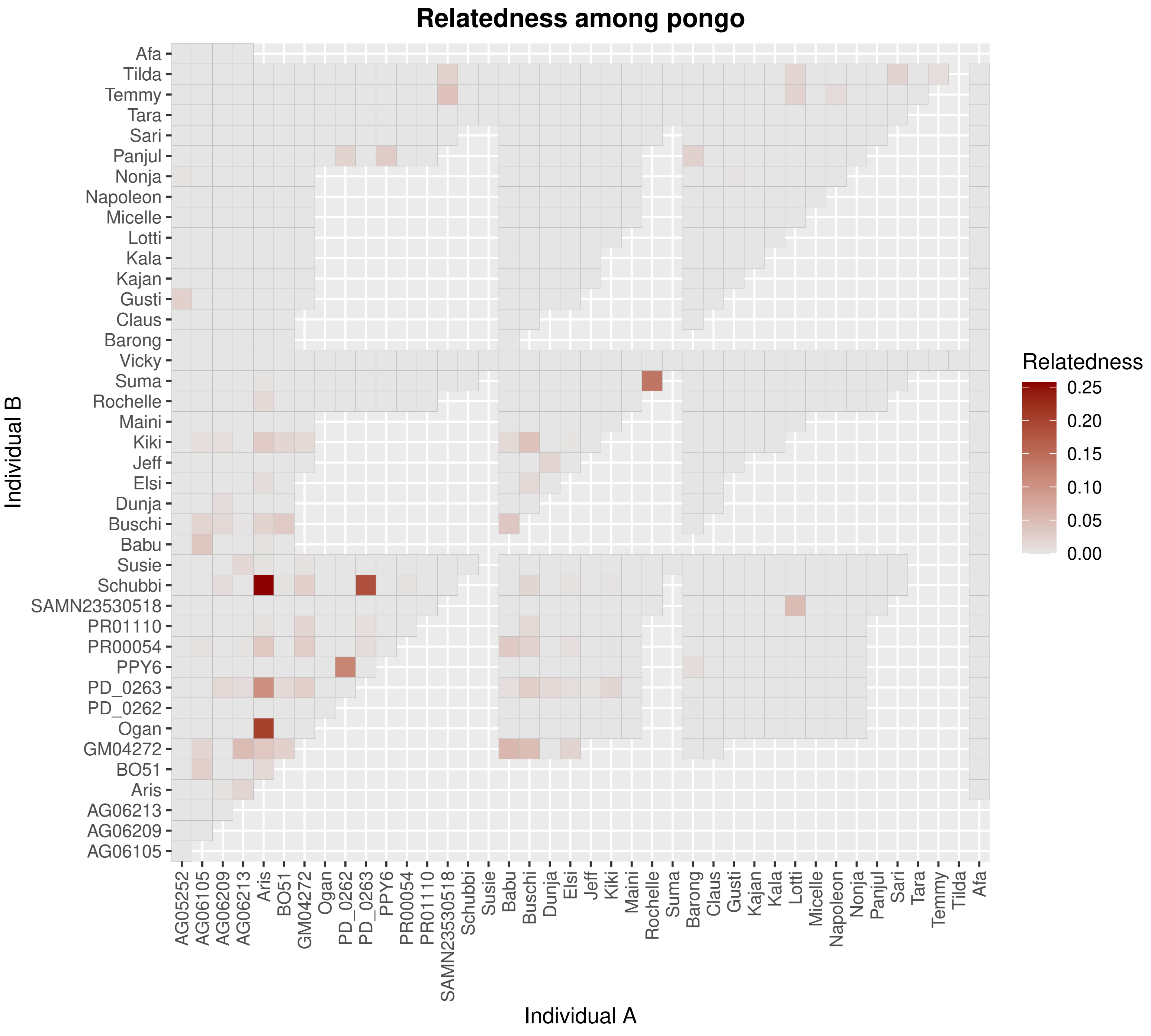

Relatedness among pongo
Individual B
Individual A
Afa
Tilda
Temmy
Tara
Sari
Panjul
Nonja
Napoleon
Micelle
Lotti
Kala
Kajan
Gusti
Claus
Barong
Vicky
Suma
Rochelle
Maini
Kiki
Jeff
Elsi
Dunja
Buschi
Babu
Susie
Schubbi
SAMN23530518
PR01110
PR00054
PPY6
PD_0263
PD_0262
Ogan
GM04272
BO51
Aris
AG06213
AG06209
AG06105
AG05252
Relatedness
0.25
0.20
0.15
0.10
0.05
0.00

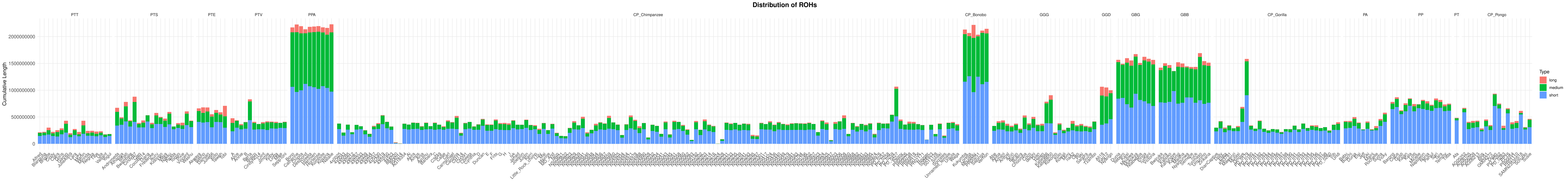
Distribution of ROHs
PTT
PTS
PTE
PTV
PPA
CP_Chimpanzee
CP_Bonobo
GGG
GGD
GBG
GBB
CP_Gorilla
PA
PP
PT
CP_Pongo
Cumulative Length
2000000000
1500000000
1000000000
500000000
0
Type
long
medium
short

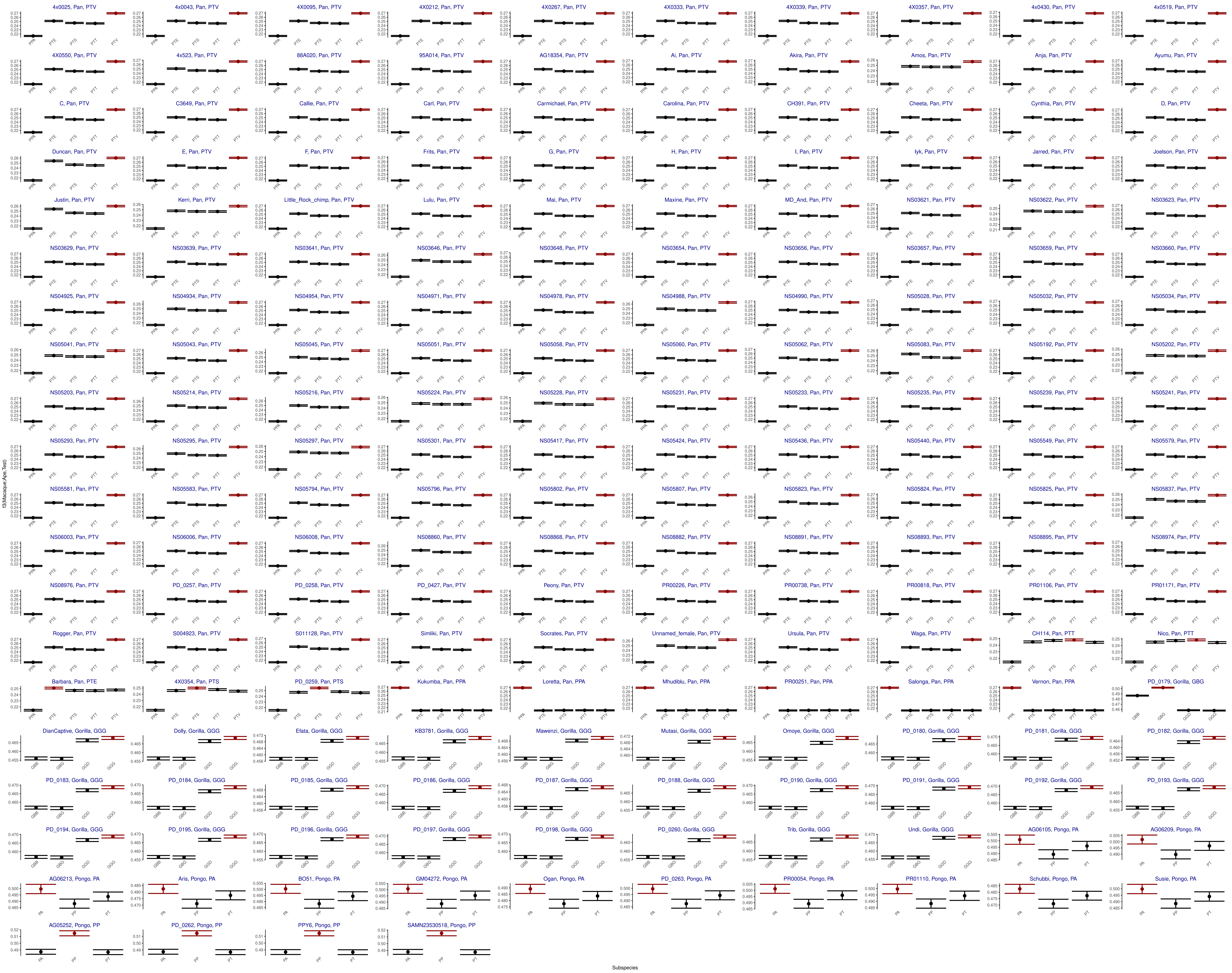
f3(Macaque;Ape,Test)
Subspecies

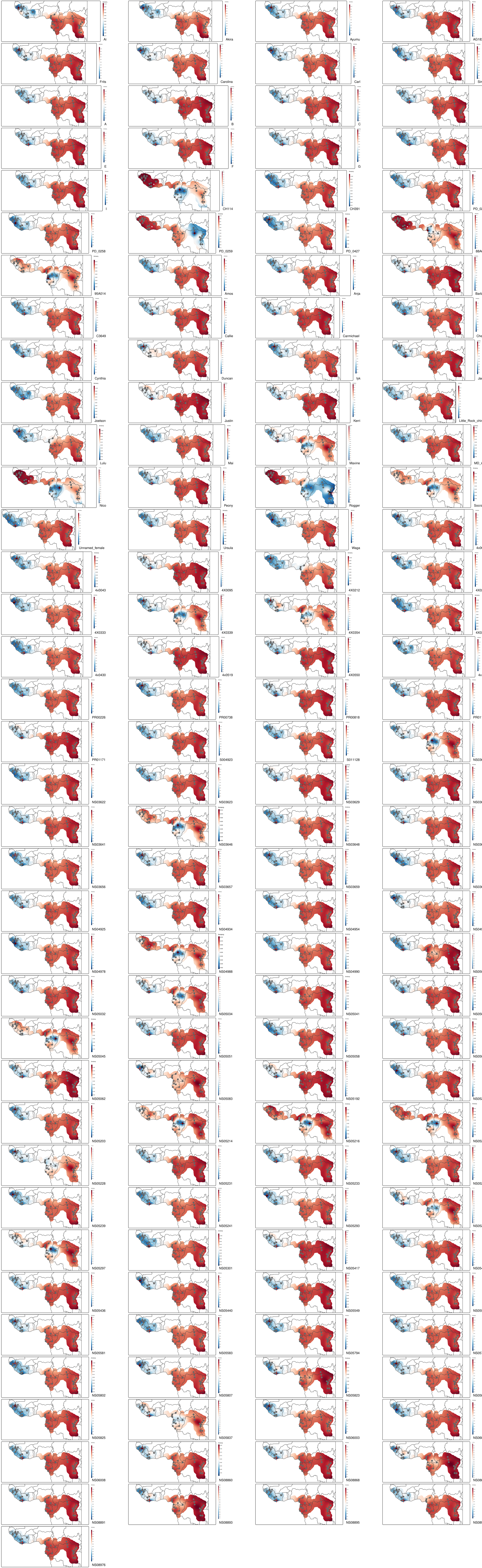

Ai
Akira
Ayumu
AG18354
Frits
Carolina
Carl
Simliki
A
B
C
D
E
F
G
H
I
CH114
CH391
PD_0257
PD_0258
PD_0259
PD_0427
88A020
95A014
Amos
Anja
Barbara
C3649
Callie
Carmichael
Cheeta
Cynthia
Duncan
Iyk
Jarred
Joelson
Justin
Kerri
Little_Rock_chimp
Lulu
Mai
Maxine
MD_And
Nico
Peony
Rogger
Socrates
Unnamed_female
Ursula
Waga
4x0025
4x0043
4X0095
4X0212
4X0267
4X0333
4X0339
4X0354
4X0357
4x0430
4x0519
4X0550
4x523
PR00226
PR00738
PR00818
PR01106
PR01171
S004923
S011128
NS03621
NS03622
NS03623
NS03629
NS03639
NS03641
NS03646
NS03648
NS03654
NS03656
NS03657
NS03659
NS03660
NS04925
NS04934
NS04954
NS04971
NS04978
NS04988
NS04990
NS05028
NS05032
NS05034
NS05041
NS05043
NS05045
NS05051
NS05058
NS05060
NS05062
NS05083
NS05192
NS05202
NS05203
NS05214
NS05216
NS05224
NS05228
NS05231
NS05233
NS05235
NS05239
NS05241
NS05293
NS05295
NS05297
NS05301
NS05417
NS05424
NS05436
NS05440
NS05549
NS05579
NS05581
NS05583
NS05794
NS05796
NS05802
NS05807
NS05823
NS05824
NS05825
NS05837
NS06003
NS06006
NS06008
NS08860
NS08868
NS08882
NS08891
NS08893
NS08895
NS08974
NS08976

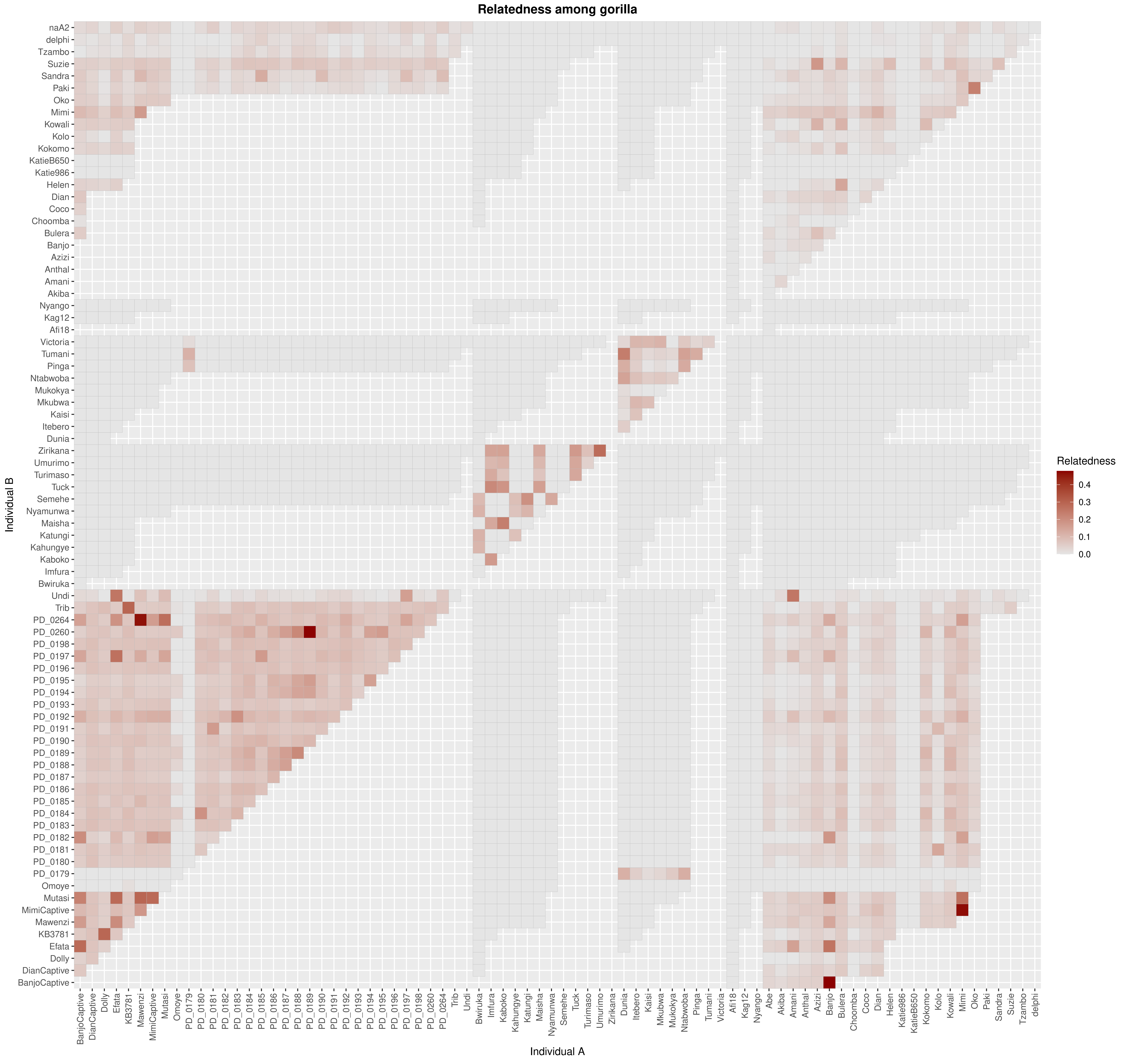

Relatedness among gorilla
Individual A
Individual B
Relatedness
0.4
0.3
0.2
0.1
0.0

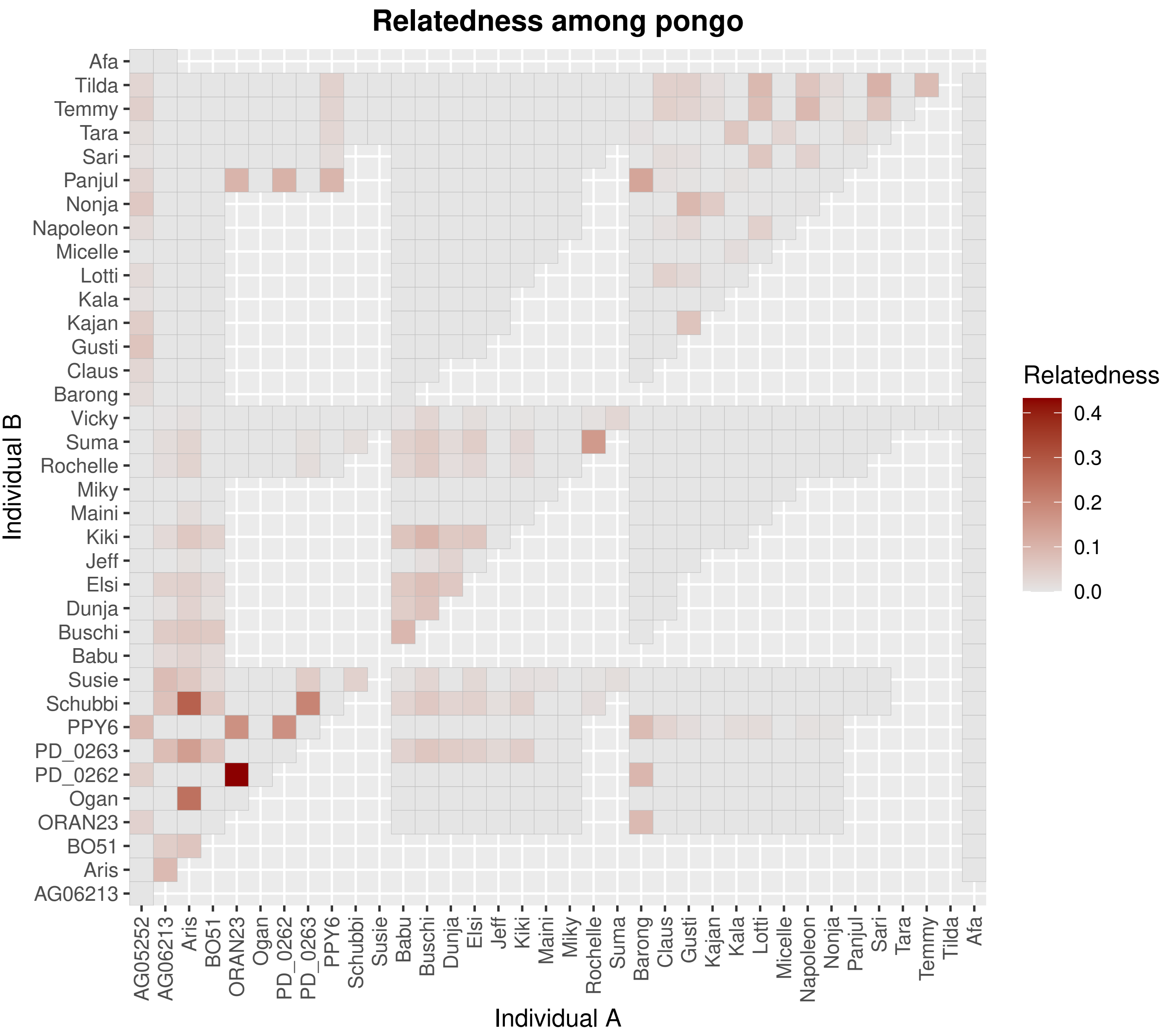
Relatedness among pongo
Individual B
Afa
Tilda
Temmy
Tara
Sari
Panjul
Nonja
Napoleon
Micelle
Lotti
Kala
Kajan
Gusti
Claus
Barong
Vicky
Suma
Rochelle
Miky
Maini
Kiki
Jeff
Elsi
Dunja
Buschi
Babu
Susie
Schubbi
PPY6
PD_0263
PD_0262
Ogan
ORAN23
BO51
Aris
AG06213
Individual A
AG05252
AG06213
Aris
BO51
ORAN23
Ogan
PD_0262
PD_0263
PPY6
Schubbi
Susie
Babu
Buschi
Dunja
Elsi
Jeff
Kiki
Maini
Miky
Rochelle
Suma
Barong
Claus
Gusti
Kajan
Kala
Lotti
Micelle
Napoleon
Nonja
Panjul
Sari
Tara
Temmy
Tilda
Afa
Relatedness
0.4
0.3
0.2
0.1
0.0

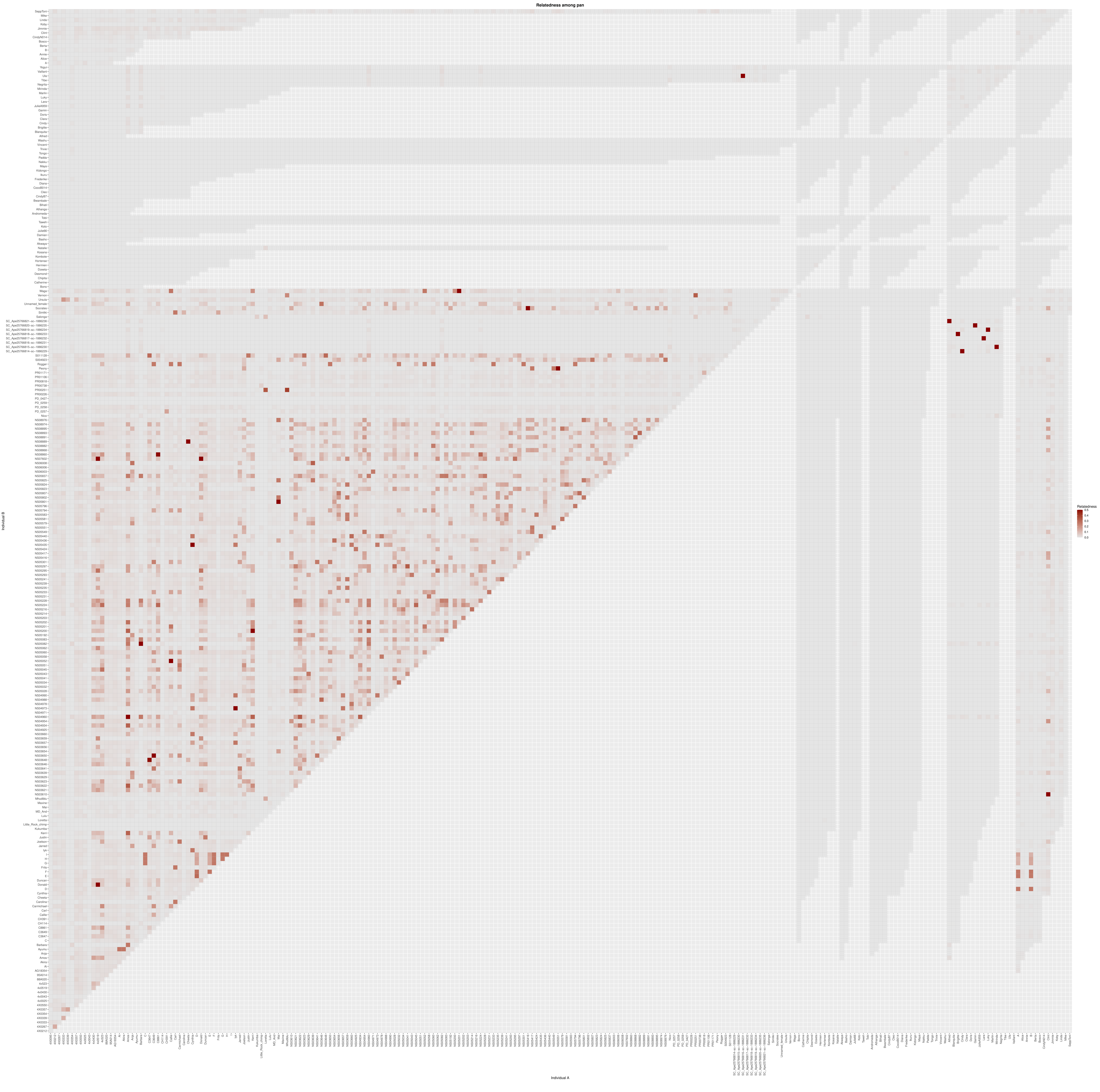
Relatedness among pan
Individual A
Individual B
Relatedness
0.5
0.4
0.3
0.2
0.1
0.0
